# The formation of intramolecular secondary structure brings mRNA ends in close proximity

**DOI:** 10.1101/289496

**Authors:** Wan-Jung C. Lai, Mohammad Kayedkhordeh, Erica V. Cornell, Elie Farah, Stanislav Bellaousov, Robert Rietmeijer, David H. Mathews, Dmitri N. Ermolenko

## Abstract

A number of protein factors regulate protein synthesis by bridging mRNA ends or untranslated regions (UTRs). Using experimental and computational approaches, we show that mRNAs from various organisms, including humans, have an intrinsic propensity to fold into structures in which the 5’ end and 3’ end are ≤ 7 nm apart irrespective of mRNA length. Computational estimates performed for ∼22,000 human transcripts indicate that the inherent proximity of the ends is a universal property of most, if not all, mRNA sequences. Only specific RNA sequences, which have low sequence complexity and are devoid of guanosines, are unstructured and exhibit end-to-end distances expected for the random coil conformation of RNA. Our results suggest that the intrinsic proximity of mRNA ends may facilitate binding of translation factors that bridge mRNA 5’ and 3’ UTRs. Furthermore, our studies provide the basis for measuring, computing and manipulating end-to-end distances and secondary structure in mRNAs in research and biotechnology.

## INRODUCTION

Regulation of mRNA translation in eukaryotes involves protein-mediated interactions between mRNA ends. Translation initiation requires the recruitment of the small ribosomal subunit to the 5’ end of the mRNA^1^. The formation of the initiation complex is stimulated by the interaction between the 5’ mRNA cap-binding protein eIF4E and the 3’ end poly(A) tail binding protein PABP, which is mediated through their binding to different parts of the translational factor eIF4G^2,3^. The eIF4E•eIF4G•PABP complex is thought to enhance translation initiation by circularizing the mRNA and forming the “closed-loop” structure^4-6^. The mechanism by which the mRNA closed loop enhances proteins synthesis is not well understood.

Remarkably, translation initiation of many eukaryotic mRNAs is also regulated by sequences in their 3’ UTRs and controlled by the formation of protein bridges between the 5’ and 3’ UTRs. For example, the 3’ UTR regulatory sequences recruit protein complexes (e.g. CPEB•Maskin, Bruno•Cup, or GAIT complex), which inhibit translation by interacting with either eIF4E or eIF4E•eIF4G bound to the 5’ end of mRNA^7^. The pervasiveness of protein bridges between mRNA UTRs in the evolution of translation regulation is puzzling because of the significant entropic cost expected for protein-mediated mRNA circularization^8^.

The entropic penalty for the formation of protein bridges between mRNA ends may be partially mitigated by mRNA compaction through intramolecular basepairing interactions. Recent theoretical analyses suggested that the 5’ and 3’ ends of long (1,000-10,000 nucleotide-long) RNAs are always brought in the proximity of few nanometers of each other regardless of RNA length and sequence because of the intrinsic propensity of RNA to form widespread intramolecular basepairing interactions^8-10^. One study predicted that the 5’ to 3’ end distance in RNAs is 3 nm, on average^8^. These theoretical predictions were tested by single-molecule Förster resonance energy transfer (smFRET) measurements of end-to-end distances in several viral RNAs and mRNAs from the fungus *Trichoderma atroviride*, which varied in length between 500 and 5,000 nucleotides and were folded *in vitro* in the absence of any protein factors^11^.

Experimentally-derived end-to-end distances in RNA molecules, in which FRET was detected, ranged between 5 and 9 nanometers^11^. However, this study did not determine the average end-to-end distance in each transcript since only molecules showing FRET were detected. Thus, it is possible that RNA molecules with 5-9 nm-long end-to-end distances account for only a minor fraction of each examined transcript.

The hypothesis that the closeness of RNA ends is a universal property of all natural transcripts remains to be systematically tested. The propensity of human mRNAs to fold into structures with short end-to-end distances has not been examined. It is unclear to what extent end-to-end distance may vary between different transcripts. Sequence features that define RNA potential to fold into structures with short end-to-end distances are unknown. Elucidating whether closeness of the 5’ and 3’ ends is an intrinsic propensity of all mRNAs may have important implications for various aspects of mRNA metabolism including translation, splicing or degradation. For example, the closeness of mRNA ends may underlie translation regulation mediated by various protein complexes that bridge mRNA UTRs.

Here, we use FRET measurements and computational analysis of RNA structure to examine the end-to-end distances in mRNAs from several species, including humans. We find that most, if not all, mRNAs have an intrinsic propensity to fold into structures with short end-to-end distances.

## RESULTS

### The 5’ end of the 5’ UTR and 3’ end of the 3’ UTR of human mRNAs are intrinsically close

We experimentally determined the end-to-end distance in a number of mRNAs using FRET between fluorophores introduced at each end of the mRNAs (**Fig. 1a**). The range of FRET sensitivity (1 to 10 nm for Cy3-Cy5 pair^12^) to distance changes matches the theoretically predicted array of distances between the 5’ and 3’ ends of structured RNAs^8,10^. We selected yeast and human mRNAs that encode abundant housekeeping proteins and have well-annotated 5’ and 3’ UTR sequences, such as yeast RPL41A (ribosomal protein L41A) and human GAPDH (glyceraldehyde-3-phosphate dehydrogenase) (**Supplementary Table 1**). In addition, we used rabbit β-globin and firefly luciferase (Fluc) mRNAs that have been used as canonical “standard” mRNAs in many previous mechanistic studies of eukaryotic translation. Selected mRNAs were less than 2,000 nucleotides long to ensure high yields of *in vitro* run-off transcription by T7 RNA polymerase **(Supplementary Table 1)**.

**Table 1.**
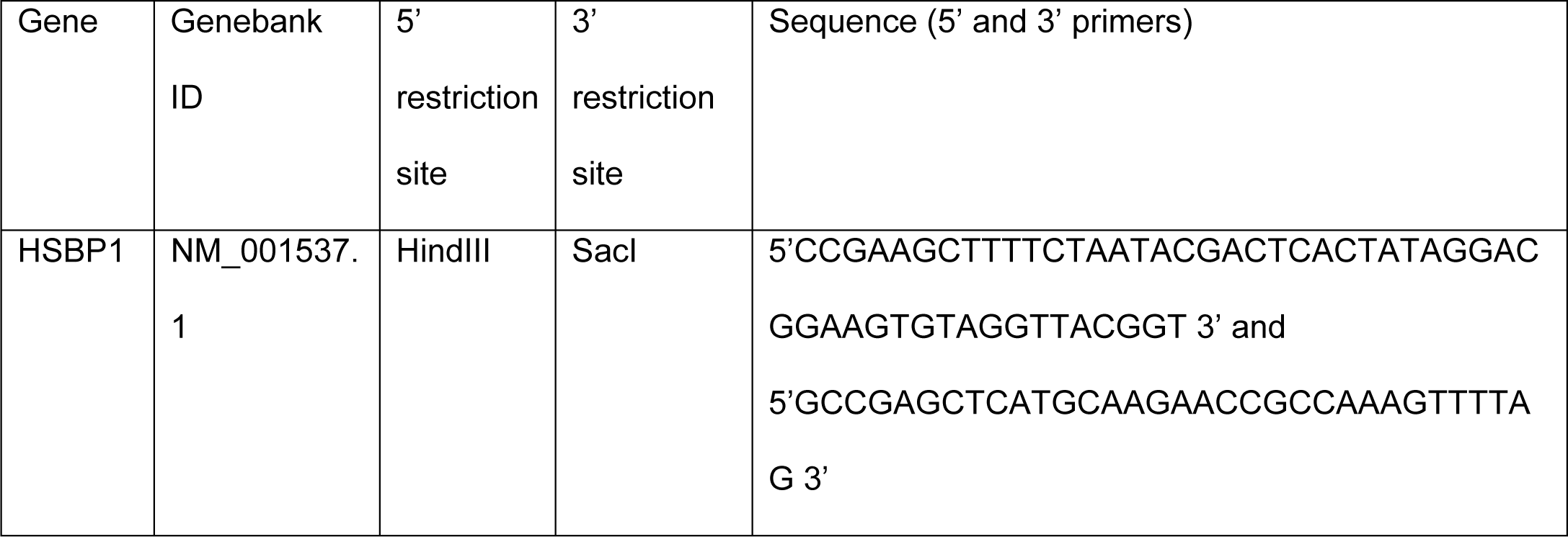
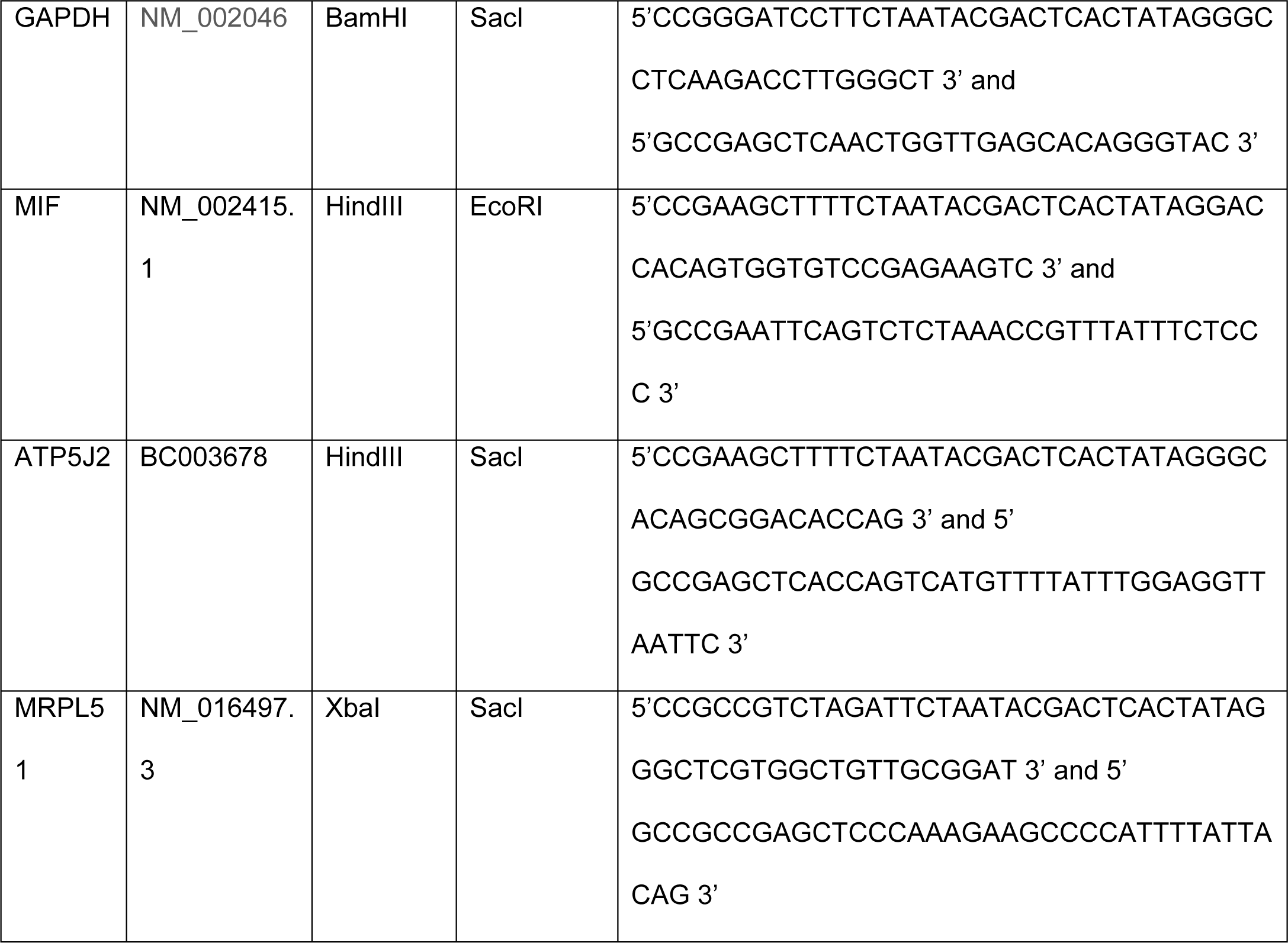
Primers used for cloning of mRNA-encoding sequences from HeLa cDNA

**Figure 1.**
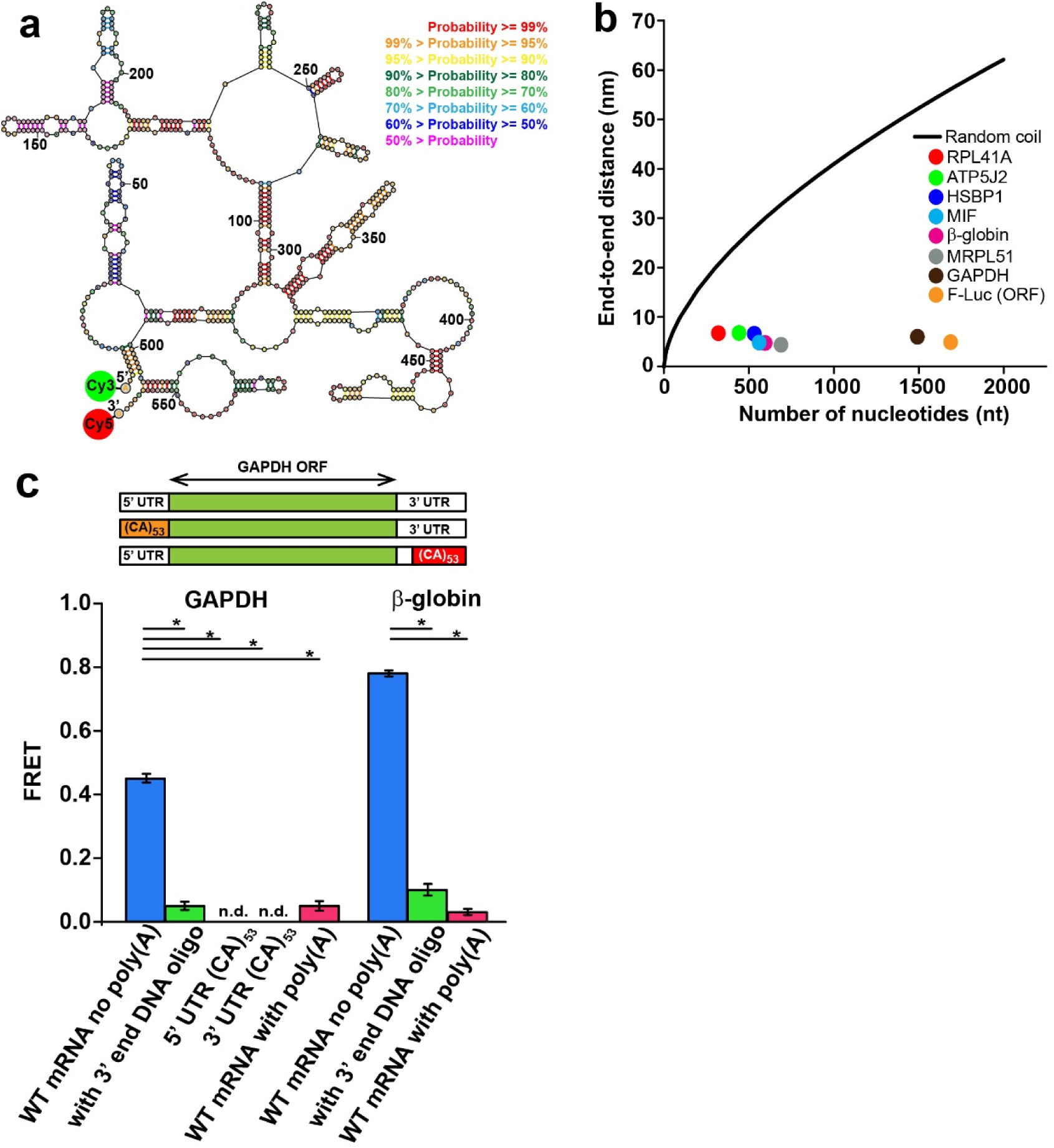
mRNA ends are brought within FRET distance via the formation of intramolecular basepairing interactions. **(a)** An exemplary secondary structure from the ensemble of structures of human MIF mRNA lacking poly(A) tail predicted by free energy minimization. Base pair probabilities, predicted with a partition function, are color-coded. In order to measure the end-to-end distance by FRET, the 5’ and 3’ ends of mRNA were conjugated with donor (Cy3) and acceptor (Cy5) fluorophores, respectively, as indicated. **(b)** Average end-to-end distances of mRNAs, which were folded in the presence of 100 mM KCl and 1 mM MgCl_2_, were determined by ensemble FRET measurements and plotted as a function of mRNA length: yeast RPL41A (red), firefly luciferase (orange), rabbit β-globin (magenta), human ATP5J2 (green), HSBP1 (indigo), MIF (blue), MRPL51 (grey), and GAPDH (brown). The black line shows theoretically predicted end-to-end distance of unstructured RNA, assuming a freely-jointed chain model. **(c)** FRET values were measured between fluorophores attached to the 5’ and 3’ ends of the following GAPDH and β-globin mRNAs: mRNA lacking poly(A) tail (blue); mRNA lacking poly(A) tail folded in the presence of a 50-nucleotide long DNA oligonucleotide complementary to the 3’ end of mRNA (green); mRNA with poly(A) tail (red). FRET could not be detected (n.d.) in GAPDH variants, which lacked poly(A) tail and contained 53 CA repeats introduced into the 5’ or 3’ UTR of GAPDH mRNA. Each FRET value represents the mean ± standard deviation (SD) of three independent experiments. A star indicates that FRET values are significantly different, as p-values determined by the Student t-test were below 0.05.

We labeled the 5’ and 3’ ends of mRNAs, which lacked the 5’ cap and poly(A) tail, with donor (Cy3) and acceptor (Cy5) fluorescent dyes, respectively. Computational prediction of RNA secondary structure suggests that all examined mRNAs can form extensive intramolecular basepairing interactions **(Fig. 1a, Supplementary Fig. 1a)**. mRNAs were refolded in the absence of protein factors and the presence of 100 mM KCl and 1 mM MgCl_2_. These ionic conditions are considered to be optimal for translation in eukaryotic *in vitro* translation systems^13,14^. Furthermore, the 1 mM concentration of Mg^2+^ used in our experiments is similar to concentrations of free (unbound) cytoplasmic Mg^2+^ in human cells (0.5-1 mM)^15^. Energy transfer between fluorophores attached to the 5’ end of the 5’ UTR and 3’ end of the 3’ UTR was detected in all eight tested mRNAs. The average end-to-end distances, which were determined for each transcript from ensemble FRET data, were in the range of 5-7 nm irrespective of mRNA length (**Fig. 1b, Supplementary Table 1**). These distances are two to ten times shorter than those predicted for unstructured RNA by the freely jointed chain model^16,17^ of polymer theory (**Fig. 1b)**.

We next tested whether mRNA ends are brought into close proximity by basepairing interactions. Refolding of human GAPDH and rabbit β-globin mRNAs in the presence of a 50 nucleotide-long DNA oligonucleotide complementary to the 3’ end of the respective mRNA led to a dramatic reduction in the efficiency of energy transfer between fluorophores attached to RNA ends. The observed decrease in FRET efficiency is presumably due to annealing of the DNA oligonucleotide to the 3’ end of the mRNA and disruption of the intramolecular secondary structure (**Fig. 1c, Supplementary Fig. 1b-c**).

To further test the effect of intramolecular secondary structure on mRNA end-to-end distance, we replaced 106 nucleotides at the 5’ end of the 116 nucleotide-long 5’ UTR of GAPDH mRNAs with 53 CA repeats, which have low basepairing potential, to create the 5’UTR(CA) _53_GAPDH mRNA variant. Likewise, 53 CA repeats were inserted at the 3’ end of the 202 nucleotide-long 3’ UTR of GAPDH mRNA in place of 106 nucleotides of the original sequence, to make the 3’UTR(CA)_53_GAPDH mRNA variant. No energy transfer between the mRNA ends was detected in either of the GAPDH mRNA variants containing CA repeats, i.e. in 5’UTR(CA)_53_GAPDH and 3’UTR(CA)_53_GAPDH mRNAs **(Fig. 1c)**. These results indicate that the 5’ and 3’ ends of wild-type GAPDH mRNA were brought within FRET distance via the formation of intramolecular basepairing interactions.

### The 3’ poly(A) tail is not involved in intramolecular basepairing interactions, which bring the ends of the 5’ and 3’ UTRs in close proximity

mRNAs in eukaryotic cells undergo 5’ capping (attachment of 7-methyl-guanosine to the 5’ end) and polyadenylation of the 3’ end. In the experiments described above, we measured the distance between the 5’ end of the 5’ UTR and the 3’ end of the 3’ UTR of model mRNAs in the absence of the 5’ cap and the poly(A) tail because neither the 5’ cap nor adenosine repeats are likely to significantly affect secondary structure of mRNAs that lack extended uridine repeats. We tested the validity of this assumption by attaching donor and acceptor fluorophores to the ends of β-globin and GAPDH mRNAs transcribed with a 30 nucleotide-long poly(A) tail. Addition of a poly(A) tail led to a significant reduction in FRET efficiency (**Fig. 1c)**, corresponding to an increase of end-to-end mRNA distance in both GAPDH and β-globin mRNAs by ∼5 nm **(Supplementary Table 1**). This value is consistent with the ∼5 nm end-to-end distance predicted for the 30 nt-long RNA segment in random-coil conformation^18^. Therefore, the poly(A) tail is unstructured and not involved in basepairing interactions with the 5’ UTR.

### mRNAs fold into a dynamic ensemble of structures

Computational predictions suggest that mRNAs fold into an ensemble of structures with comparable thermodynamic stabilities rather than a single structure. To test this prediction, we examined end-to-end distance in individual GAPDH, β-globin and MIF mRNA molecules by measuring single-molecule (sm)FRET using total internal reflection fluorescence (TIRF) microscopy. smFRET reveals the structural dynamics of individual molecules that are masked in ensemble (bulk) FRET measurements because of signal averaging in the heterogeneous and non-synchronized population^12^. To immobilize mRNAs to the surface of the microscope slide, a 20-nucleotide-long DNA oligonucleotide conjugated to biotin was annealed in the middle of the RNA where it was computationally predicted to have a minimal effect on the overall secondary structure and end-to-end distance of the RNA^19^ (**Supplementary Fig. 1b-c**). No significant decrease in energy transfer in ensemble FRET experiments was observed when Cy3/Cy5-labeled GAPDH, β-globin and MIF mRNAs lacking poly(A) tail were folded in the presence of biotin-labeled DNA oligonucleotides (**Supplementary Fig. 2**). Hence, annealing of biotin-labeled DNA oligonucleotides did not affect end-to-end distance nor disrupt the overall RNA structure.

**Figure 2.**
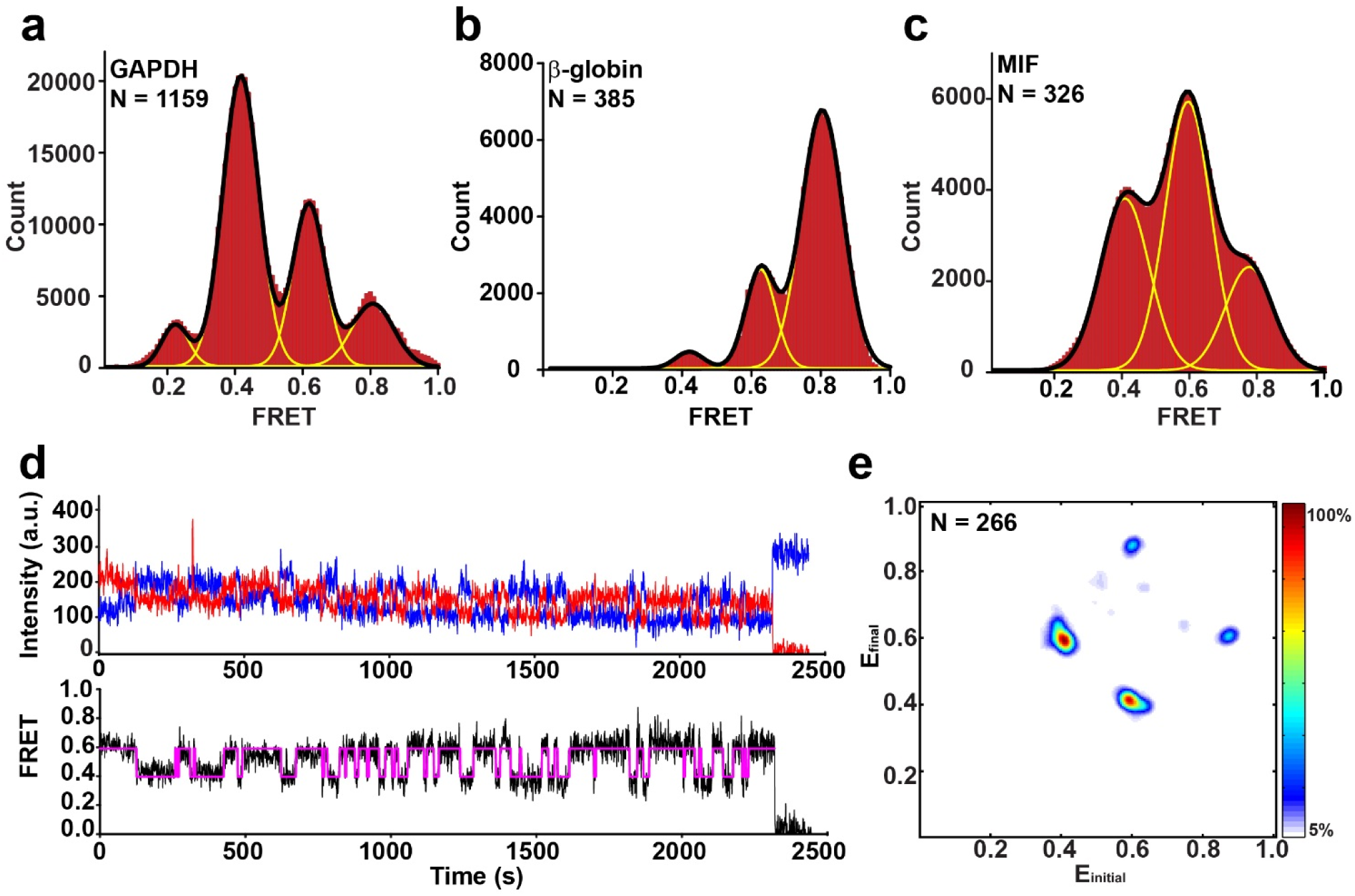
mRNAs fold into a dynamic ensemble of structures. smFRET was measured between the dyes attached to the 5’ end of the 5’ UTR and 3’ end of the 3’ UTR in GAPDH, β-globin, and MIF mRNAs. **(a-c)** Histograms, compiled from hundreds of smFRET traces, showing the distribution of the FRET values in **(a)** GAPDH mRNA, **(b)** β-globin mRNA, and **(c)** MIF mRNA folded in the presence of 100 mM KCl and 1 mM MgCl_2_. Yellow lines represent individual Gaussian fits and black lines indicate the sum of Gaussians. N is the number of single-molecule traces compiled. **(d)** Representative smFRET trace for GAPDH mRNA showing fluctuations between the 0.4 and 0.6 FRET states. Observed intensities of donor and acceptor fluorescence and the calculated apparent FRET efficiency are shown in blue, red, and black, respectively. Hidden Markov Model fit is shown in magenta. **(e)** Transition density plot (TDP) analysis of 5,114 fluctuations between different FRET states in 266 HMM-idealized FRET traces obtained for GAPDH mRNA. The frequency of transitions from the starting FRET value (x axis) to the ending FRET value (y axis) is represented by a heat map. The range of FRET efficiencies from 0 to 1 was separated into 200 bins. The resulting heat map was normalized to the most populated bin in the plot; the lower- and upper-bound thresholds were set to 5% and 100% of the most populated bin, respectively.

mRNAs were tethered to the surface of microscope slides coated with BSA-biotin/ neutravidin and then imaged by exciting the donor (Cy3) fluorescence with the green (532 nm) laser. smFRET traces acquired for GAPDH mRNA, β-globin and MIF mRNAs exhibited single-step photobleaching of both donor and acceptor fluorophores, indicating that we observe intramolecular rather than intermolecular energy transfer between mRNA ends (**Supplementary Fig. 3**). Because efficiencies of labeling of the 5’ end with Cy5 (100%) and the 3’ end of mRNA with Cy3 (∼20-30%) markedly differed, we also imaged mRNAs by exciting the acceptor (Cy5) fluorescence with the red (642 nm) laser. Single-step Cy5 photobleaching was observed in 97-99% of single-molecule traces of GAPDH, β-globin and MIF mRNAs, indicating that mRNA dimers or higher-order oligomers were essentially absent.

**Figure 3.**
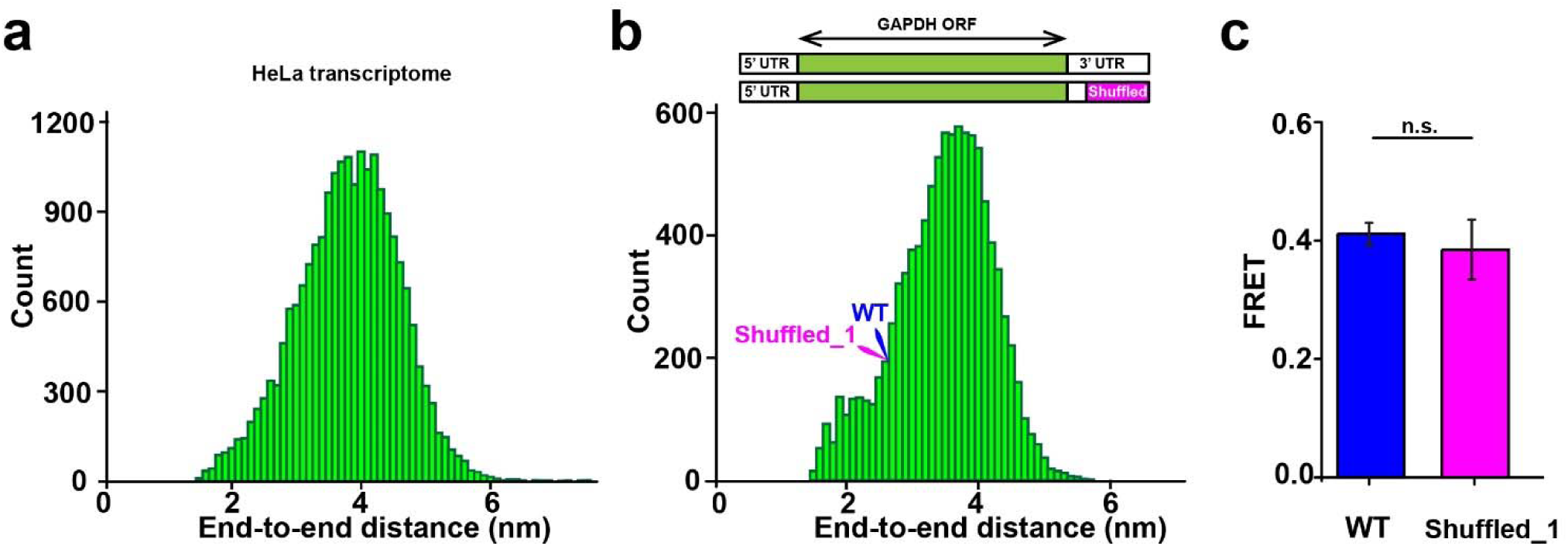
Ends of human mRNAs are universally close. **(a)** Distribution of computationally predicted average end-to-end distances in ∼22,000 mRNA sequences from the HeLa cell transcriptome. **(b)** Distribution of average end-to-end distances in 10,000 GAPDH mRNA variants, in which 106 3’ terminal nucleotides were shuffled. Predicted end-to-end distances in wild-type and shuffled_1 GAPDH mRNAs are indicated by blue and pink arrows. **(c)** Ensemble FRET values measured in wild-type (blue) and shuffled_1 (pink) GAPDH mRNA variants. Each FRET value represents the mean ± SD of three independent experiments. The difference between FRET values was not statistically significant (n.s.) as determined by the Student t-test with α of 0.05.

FRET distribution histograms, which were constructed by compiling 300-1,200 smFRET traces, are best fit to a sum of four (GAPDH mRNA) or three (β-globin and MIF mRNAs) Gaussians (**Fig. 2a-c**). The distinct FRET peaks in distribution histograms correspond to different FRET states and, thus, different mRNA end-to-end distances. During run-off transcription, T7 RNA polymerase can add one, two or three non-templated nucleotides to the 3’ RNA end in a fraction of the transcripts. To test whether the presence of multiple FRET states in FRET histograms corresponds to sequence or secondary structure heterogeneity, we varied the concentration of Mg^2+^, which is known to stabilize the secondary and tertiary structure of RNA. An increase in MgCl_2_ concentration from 0 to 2 mM during mRNA refolding reduced the fraction of molecules exhibiting lower FRET states and increased the fraction of molecules showing higher FRET values without substantially altering the positions of the peaks of the FRET states (**Supplementary Fig. 4**) in GAPDH, β-globin and MIF mRNAs. Hence, most, if not all, distinct FRET peaks of smFRET distribution histograms correspond to different interconverting structural states. Nevertheless, a contribution of sequence heterogeneity to the breadth of smFRET histograms cannot be completely ruled out.

**Figure 4.**
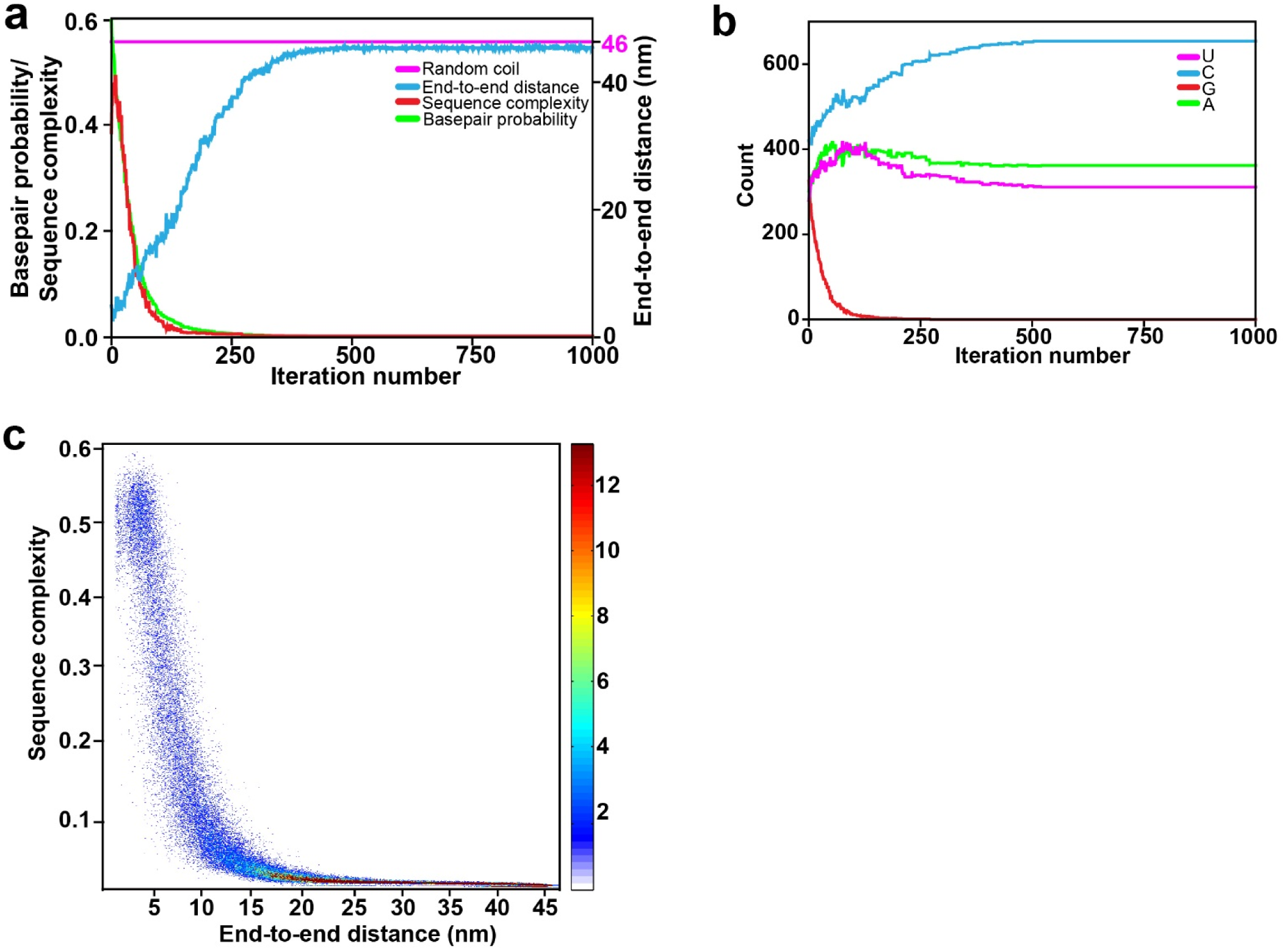
Sequence features of intrinsically-unstructured RNA sequences. The entire 1327-nt long GAPDH mRNA sequence was evolved *in silico* by a genetic algorithm to minimize average basepairing probability and produce intrinsically unstructured sequences. **(a)** End-to-end distance (blue), sequence complexity (red), and mean basepair probability (green) as functions of iteration number are shown for a single representative *in silico* sequence evolution experiment. The distance predicted for a 1327-nt long RNA in a random coil conformation is shown by the magenta line. **(b)** Evolution of nucleotide composition in a single representative *in silico* sequence transformation experiment shown in **(a)**. Frequency of adenosine (A), cytidine (C), guanosine (G), and uridine (U) are shown in magenta, blue, red, and green, respectively. **(c)** Surface contour plots generated from 500 independent *in silico* sequence evolution experiments show changes of sequence complexity (y-axis) as a function of end-to-end distance (x-axis). The range of sequence complexity from 0 to 0.6 was separated into 2,000 bins. The range of end-to-end distance from 1.6 to 46 nm was separated into 500 bins. The resulting heat map shows the frequency count.

An increase in MgCl_2_ concentration during RNA folding also raised ensemble FRET values (**Supplementary Fig. 5**), demonstrating consistency between ensemble and single-molecule experiments. Interestingly, FRET between the 5’ and 3’ ends in β-globin and MIF mRNAs showed stronger dependence on Mg^2+^ concentration than FRET in GAPDH mRNA (**Supplementary Fig. 4 and 5**), suggesting that in addition to the formation of secondary structure, intramolecular tertiary interactions may be involved in bringing the ends of β-globin and MIF mRNAs in close proximity.

**Figure 5.**
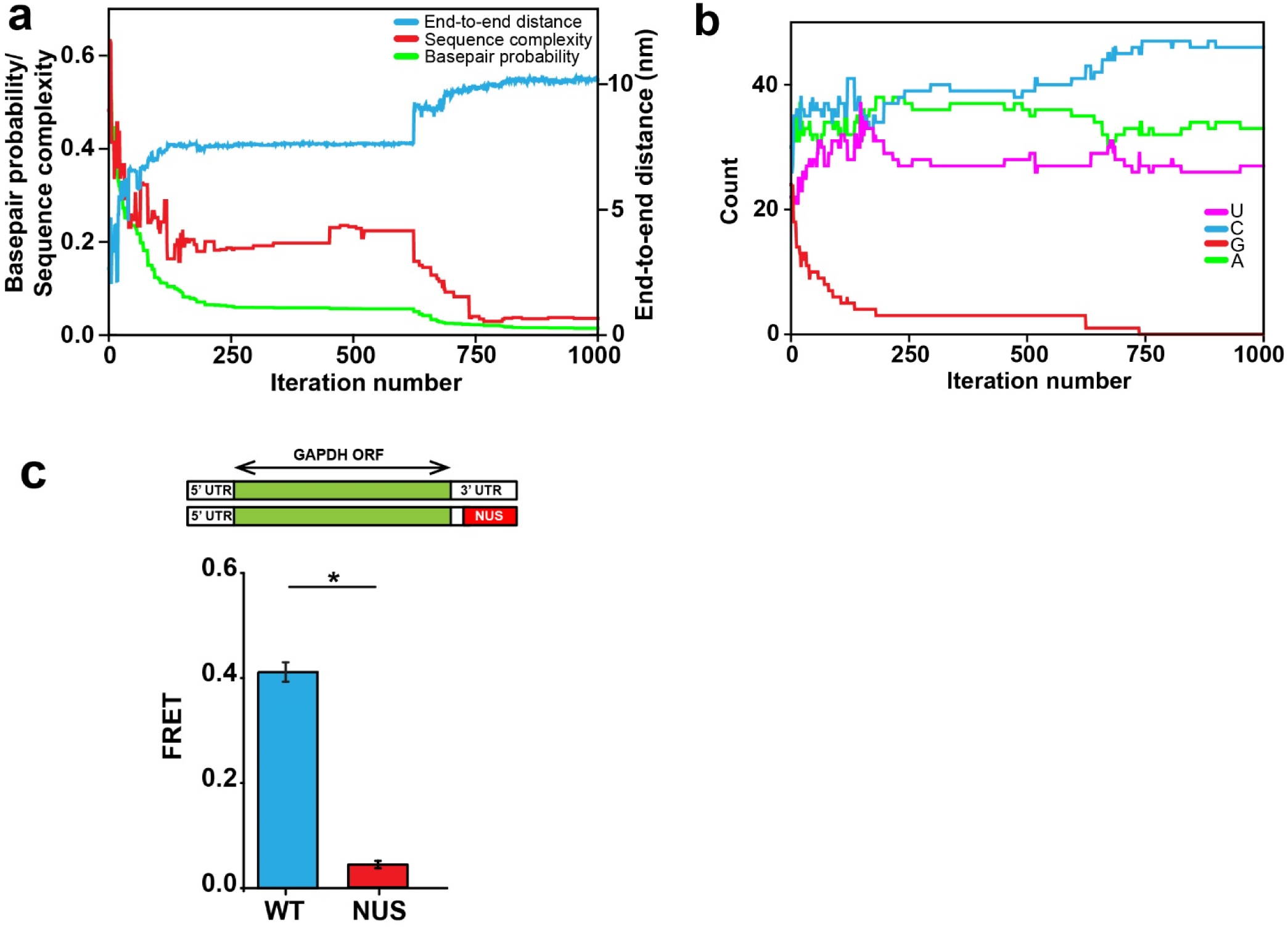
Manipulation of end-to-end distances in GAPDH mRNAs *in silico* with a genetic algorithm. 106 nucleotides in the 3’ end of the 3’ UTR in GAPDH mRNA are computationally evolved *in silico* by a genetic algorithm. **(a)** End-to-end distance (blue), sequence complexity (red), and basepair probability (green) as functions of iteration number are shown for a single *in silico* sequence evolution experiment. **(b)** Evolution of nucleotide composition in the sequence evolution *in silico* experiment shown in **(a)**. Frequency of adenosine (A), cytidine (C), guanosine (G), and uridine (U) are shown in magenta, blue, red, and green, respectively. **(c)** FRET values were measured in wild-type GAPDH mRNA (blue) and the GAPDH mRNA variant with the non-repetitive unstructured (NUS) 106 nucleotide sequence in the 3’ end of the 3’ UTR (red) designed by a genetic algorithm. Each FRET value represents the mean ± SD of three independent experiments. A star indicates that FRET values are different, as p-values determined by the Student t-test were below 0.05.

Consistent with the idea that mRNAs fold into a dynamic ensemble of several structural states with multiple end-to-end distances, individual smFRET traces in GAPDH, β-globin and MIF mRNAs showed spontaneous fluctuations between distinct FRET states (**Fig. 2d, Supplementary Fig. 3**). Using GAPDH mRNA as an example, we further explored the statistics of fluctuations between FRET states via Hidden Markov Model (HHM) and Transition Density Plot analyses^20,21^. Consistent with FRET distribution histograms, individual GAPDH mRNA molecules predominantly fluctuated between ∼0.4, 0.6 and 0.8 FRET states at frequencies of ∼0.1 - 0.03 s^-1^ (**Fig. 2e, Supplementary Table 2**). These rates are similar to previously measured kinetics of the spontaneous transition between two alternative 5 basepair-long RNA helixes^22^, indicating that FRET changes observed in GAPDH mRNA may correspond to analogous structural rearrangement. Although a few (1%) of the dynamic smFRET traces showed fluctuations between three (0.2, 0.4 and 0.8) or all four (0.2, 0.4, 0.6 and 0.8) FRET states (**Supplementary Fig. 3**), the majority (99%) of traces showed reversible fluctuations that transitioned between just two states (either between 0.4 and 0.6 or between 0.6 and 0.8; **Fig. 2e**). These data suggest there may be two dynamic structurally-distinct sub-populations of GAPDH mRNA.

**Table 2.**
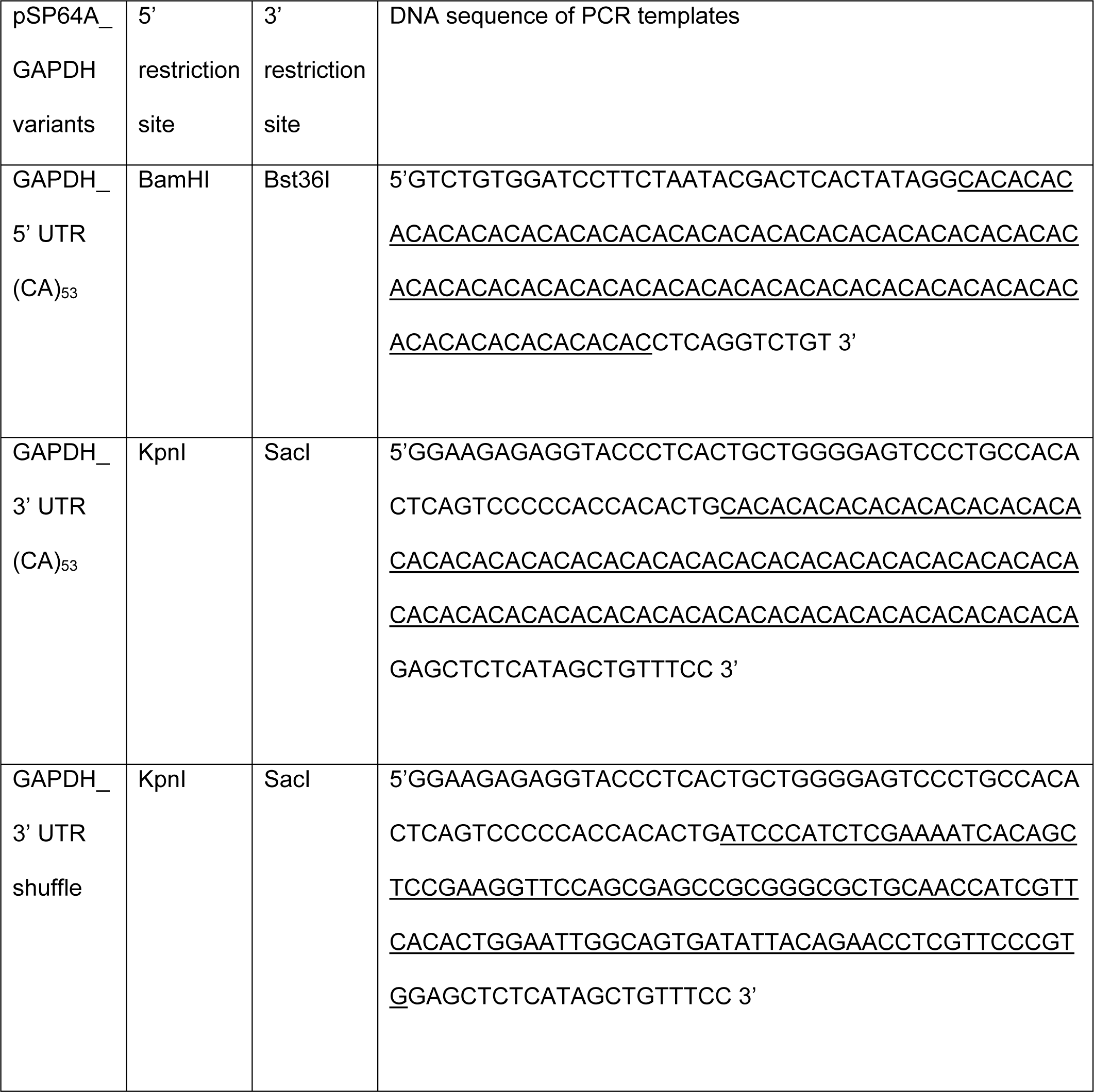
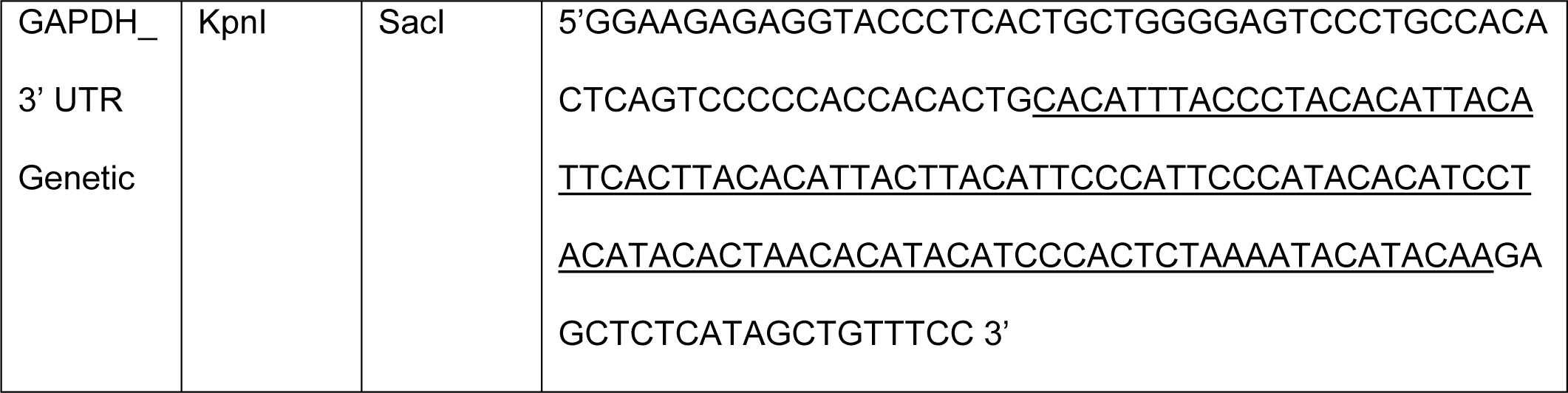
PCR templates for cloning unstructured or randomized sequences (underlined) into the UTRs of GAPDH mRNA.

### Algorithm for the prediction of end-to-end distances in natural mRNAs through computation

Using the RNAstructure software package^23^ and a freely jointed chain polymer theory^18^, we developed a new algorithm for modeling the distribution of end-to-end distances for the folding ensemble in natural mRNAs to test the hypothesis about the proximity of mRNA ends at a transcriptome-wide level. In this algorithm, a representative thermodynamic ensemble of structures is selected by stochastic sampling^24^, and then the distance between the 5’ and 3’ ends is estimated for each member of the sample in nanometers. The calculation employs two segment sizes (unpaired nucleotides and helix ends), which are estimated based on the freely jointed chain model of polymer theory^16-18^. We do not consider the presence of the poly(A) tail of the mRNA because the poly(A) tail is unlikely to make base-pairing interactions with the rest of the mRNA (**Fig. 1c**). Thus, we estimate the distance between the 5’ end of mRNA and the 3’ end of the 3’ UTR at the junction with poly(A) tail. Our algorithm generates a histogram of estimated distances and, thus, examines both average end-to-end distance and the distribution of end-to-end distances in the population of RNA structures.

Average end-to-end distances derived from our ensemble FRET measurements correlate reasonably well with distances predicted for the same mRNAs using computation with a linear regression coefficient, r^2^, of 0.67 (**Supplementary Table 1, Supplementary Fig. 6**). Deviations between predicted and experimentally measured end-to-end distances do not exceed 3 nm and, at least in part, may result from perturbations of fluorescent properties of the donor and acceptor fluorophores due to local environmental effects, which may lead to a 0.5-1 nm error in determination of FRET-derived distances^12,25^. Furthermore, FRET might overestimate the average end-to-end distance because a fraction of mRNA may be misfolded or unfolded under chosen experimental conditions. Hence, our computational algorithm adequately predicts the end-to-end distance in the ensemble of folded RNA molecules and can be used to examine end-to-end distances in the human transcriptome.

### The inherent closeness of the ends is a universal property of most, if not all, human mRNA sequences

We used our algorithm to predict the end-to-end distance in ∼22,000 transcripts of the HeLa human cell transcriptome. The predicted end-to-end distances were relatively narrowly distributed with a population mean of ∼ 4 nm (**Fig. 3a**). Hence, the propensity of folding into structures with short end-to-end distances is common to all human mRNAs. Furthermore, closeness of mRNA ends appears to be largely independent of nucleotide sequence and mRNA length.

To further explore the dependence of the end-to-end distance on RNA sequence, we estimated end-to-end distances in 10,000 variants of GAPDH mRNA, in which a segment of 106 nucleotides at 3’ end of the 3’ UTR was shuffled while preserving the original adenosine/guanosine/cytosine/uracil ratio (**Fig. 3b**). Similar to the distribution of end-to-end distances in the HeLa cell transcriptome, end-to-end distances in GAPDH variants with a shuffled sequence in the 3’ UTR were narrowly distributed with a population mean of ∼3.8 nm (**Fig. 3b**). To experimentally test these computational estimates, we cloned, transcribed without the 3’ poly(A) tail and labeled with Cy3/Cy5 fluorophores one of shuffled GAPDH variants (“Shuffled_1”). End-to-end distances for the original wild-type and Shuffled_1 GAPDH mRNAs were predicted to have equal end-to-end distances. Consistent with computational prediction, ensemble FRET values measured in wild-type and Shuffled_1 GAPDH mRNA variants were essentially indistinguishable (**Fig. 3c, Supplementary Table 1**). This result provides additional evidence that mRNA end-to-end distance is largely sequence independent. Hence, random RNA sequences tend to form secondary structure.

### Unstructured RNAs are devoid of guanosines and have low sequence complexity

Although we find that the ends of most RNA sequences are inherently close, we have also demonstrated that the introduction of CA-repeats, which are known to have low basepairing potential, increase end-to-end distance in RNA. To further investigate the relationships between sequence properties, basepairing potential and end-to-end distance of RNA, we evolved the human GAPDH mRNA sequence *in silico* using a genetic algorithm. In this newly-developed algorithm, populations of sequences are evolved either by random mutation or by crossover (combination of two sequences from the population), and then sequences with the lowest mean basepairing probabilities are selected for subsequent iterative refinement (details can be found in **Methods**). We performed 500 independent *in silico* evolution transformations of the GAPDH mRNA sequence; each of these transformations entailed 1,000 mutation/sequence selection rounds. For each iteration of sequence evolution, we estimated RNA end-to-end distance and sequence linguistic complexity^26,27^. This quantity is bounded to be greater than zero and less than or equal to 1, where complexities reflect a larger diversity in oligonucleotide sequences within a sequence (the quantitative definition can be found in **Methods**).

*In silico* evolution transformations of the GAPDH mRNA sequence revealed that a reduction in average basepairing probability leads to an increase in end-to-end distance of RNA (**Fig. 4a**). In addition, the RNA sequence become enriched with cytosines and depleted of guanosines (**Fig. 4b**). Guanosines are likely depleted because, in addition to Watson-Crick G-C base pairs, they can form wobble base pairs with uracils. G-U wobble base pairs have comparable thermodynamic stability to Watson–Crick A-U base pairs and are nearly isosteric to them^28,29^.

Reduction in average basepairing probability is also accompanied by the decrease in sequence linguistic complexity (**Fig. 4a**) leading to the emergence of degenerate and repetitive sequences. The heat map showing alterations in the distribution of sequence complexities and end-to-end distances over 500 independent *in silico* evolution transformations of the GAPDH mRNA sequence indicates that the sequence complexity and end-to-end distance undergo anti-correlated changes (**Fig. 4c**).

Although the decrease in sequence complexity levels off and sequences become completely depleted of guanosines after ∼200 iterations, basepairing probability and end-to-end distance continue evolving until they plateau after ∼500 iterations. The maximal value of end-to-end distance (46 nm) achieved during *in silico* sequence evolution of the 1327 nt-long GAPDH mRNA is equal to the end-to-end distance predicted for RNA of this length in the random-coil conformation. Therefore, guanosine depletion and diminishing of sequence complexity are necessary but not sufficient to convert structured RNA into a completely unstructured conformation. Only specific, low-complexity sequences of adenosines, cytosines and uracils adopt the random coil conformation. Hence, if intrinsically unstructured RNA sequences occur in organisms, then they likely play an important biological role and emerged as results of intense natural selection.

### Rational design of non-repetitive sequences with low basepairing potential and long end-to-end distances

We and others find that long repetitive sequences, such as CA and CAA repeats, which are commonly introduced into RNA to disrupt RNA secondary structure, are notoriously difficult to maintain and propagate in live cells^30^. To overcome this problem, we employed our genetic algorithm to generate 500 sequences of human GAPDH mRNA, in which the last 106 nucleotides of the 3’ UTR were evolved into non-repetitive, intrinsically-unstructured sequences (**Fig. 5a-b**). During sequence evolution, the selection criteria were changed to consider both linguistic complexity and mean basepairing probability to increase end-to-end distance and also avoid highly-repetitive sequences. One of the resulting GAPDH mRNA sequences was cloned, transcribed without 3’ poly(A) tail and fluorescently labeled for FRET measurements of end-to-end distance. Consistent with computational prediction, introduction of the non-repetitive, unstructured sequence into the 3’ end of the 3’ UTR of GAPDH mRNA led to a dramatic decrease in energy transfer between fluorophores attached to mRNA ends (**Fig. 5c**). This proof-of-principle experiment demonstrates that our new genetic algorithm for sequence evolution can help to design non-repetitive unstructured RNA sequences, which may be employed to study roles of RNA secondary structure in different aspects of RNA function.

## DISCUSSION

Taken together, our data strongly support the hypothesis^8-10^ that RNA as a polymer has an intrinsic propensity to fold into structures in which the 5’ and 3’ ends are just a few nm apart. Furthermore, we show that the ends of natural human mRNA sequences, folded in the absence of protein factors, are universally close. This occurs not only because of base pairs between nucleotides in the 5’ and 3’ UTRs but also because stem loop formation across whole sequences tends to shorten the end-to-end distance (**Fig. 1a, Supplementary Fig. 1**). Our computation and smFRET studies also show that each mRNA sequence folds into a dynamic ensemble of structures with distinct but nevertheless short end-to-end distances. Hence, in the ensemble of structures, mRNA ends are brought in close proximity by a number of alternative helixes formed between the 5’ and 3’ UTRs rather than by one specific set of base pairs between mRNA ends.

At least to some degree, the intrinsic mRNA propensity of folding into structures with short end-to-end distances is likely realized in live cells. *In vivo*, mRNA secondary structure may be disrupted by RNA binding proteins and RNA helicases^31,32^. Furthermore, the ribosome efficiently unwinds the secondary structure of mRNA within an Open Reading Frame (ORF) by translocating along the mRNA during the elongation of the polypeptide chain^33,34^. However, mRNA interactions with the ribosome and RNA-binding proteins are dynamic and, thus, at least transiently, mRNA may fold *in vivo*. Indeed, several recent transcriptome-wide chemical probing studies in yeast, plant and human cells showed that mRNAs adopt extensive intramolecular secondary structure *in vivo*^31,35-38^. A number of structured elements within the mRNA, including bacterial riboswitches^39^, frameshift-inducing hairpins and pseudoknots of eukaryotic viruses^40^, Internal Ribosome Entry Sites (IRES)^41^, Iron Response Elements (IRE) in the 5’ UTR of transcripts coding for proteins involved in iron metabolism,^42^ and Cap-Independent Translational Enhancers (CITEs)^43^, were shown to regulate translation initiation. Studies have also shown that accessibility of protein binding sites on mRNA and mRNA polyadenylation are also governed by RNA secondary structure^32,44,45^. Hence, at least to some extent, the secondary structure of mRNA can be maintained in the cell despite the presence of RNA helicases and other RNA-binding proteins^32,36-38^.

Published data also provide several lines of evidence that, in live cells, the 3’ end of the 3’ UTR may be brought near the 5’ end of mRNA through intramolecular basepairing interactions within mRNA rather than by the protein-mediated mRNA circularization. For example, eIF4E•eIF4G was shown to crosslink to the 3’ end of the 3’UTR of yeast transcripts in live cells in the absence of PABP^46^, which is thought to be required to circularize mRNA into the closed-loop structure. In addition, *in vivo* mapping of RNA-RNA interactions using psoralen cross-linking in yeast and human cells detected intramolecular basepairing interactions between distant segments of mRNA, including interactions between the 5’ and 3’ UTRs^36-38^.

The inherent closeness of mRNA ends may have important implications for many aspects of mRNA biology, including translation. Indeed, a number of protein complexes that regulate translation initiation, such as eIF4E•eIF4G•PABP, simultaneously interact with the mRNA 5’ and 3’ ends or UTRs. These protein complexes may have emerged during evolution to exploit the intrinsic closeness of mRNA ends. Thus, mRNA secondary structure, which brings mRNA ends in close proximity, may stabilize binding of translation factors bridging mRNA ends.

Spontaneous fluctuations of mRNA between different structural states, which were observed in our smFRET experiments, might also play a role in translation. The presence of stable secondary structure near the 5’ mRNA cap and the start codon was previously shown to inhibit translation initiation^47,48^. It is possible that during translation initiation, the 5’ end of the 5’ UTR undergoes partial unfolding while the rest of mRNA remains folded and compact, enabling recruitment of the small ribosomal subunit to the 5’ mRNA cap.

In the course of this work, we developed approaches and tools for measuring, computing, and manipulating mRNA end-to-end distances and the secondary structure of mRNA UTRs. This methodology can now be utilized to study unknown roles of mRNA end-to-end distance and secondary structure in mRNA UTRs in protein synthesis and other facets of mRNA biology.

## METHODS

### Experimental procedures

#### Cloning of mRNA-encoding sequences

Human cDNA was used to clone mRNA-encoding sequences. To prepare human cDNA, total RNA was first extracted from HeLa cells using TRIzol® reagent (Invitrogen Life Technologies) according to the manufacturer’s protocol. Genomic DNA was removed from the sample by DNase treatment (NEB). RT-PCR was performed to synthesize cDNA using 5 μg of RNA, SuperScript III Reverse Transcriptase (Invitrogen Life Technologies) and oligo dT_23_ (Sigma), following the manufacturers’ protocols. The target genes were amplified by PCR using Q5 DNA polymerase (NEB) and 5’ and 3’ primers listed in Table 1. All 5’ primers contain a restriction site for cloning, a T7 promoter sequence (5’ TTCTAATACGACTCACTATAGG 3’), and the sequence complementary to the 5’ end of the 5’ UTR. All 3’ primers contain the sequence complementary to the 3’ end of the 3’ UTR and a restriction site for cloning.

Primers were designed based on general guidelines using the IDT *OligoAnalyzer* tool. The optimal annealing temperature for each primer pair was determined using the *NEB Tm Calculator* to set the PCR conditions. A 30 second annealing step 3°C above the Tm was used after the initial denaturation step of 30 seconds at 95°C. The extension temperature was set to 72°C for 1 min per kb. The PCR products were cloned into polylinker sites of the pSP64A vector (Promega), where the 3’ cloning site has a stretch of 30 dA:dT residues inserted between the SacI and EcoRI restriction sites. Linearization of the recombinant plasmid with EcoRI was performed for the use of run-off *in vitro* transcription by T7 RNA polymerase to prepare RNA with a synthetic 30-nt poly(A) tail.

The rabbit β-globin gene was amplified directly from rabbit globin mRNA purchased from Sigma using the forward (5’ TCGAGTAAGCTTACACTTGCTTTTGACACAACTGTG 3’) and reverse (5’ CATACAGAGCTCGCAATGAAAATAAATTTCCTTTATTAGC 3’) primers and cloned into the pSP64A vector using HindIII and SacI. Because an EcoRI restriction site is present in the sequence encoding β-globin mRNA, the second EcoRI site positioned downstream of the poly(A) track was mutated into an AgeI restriction site via the QuikChange Site-Directed Mutagenesis System (Stratagene) using internal mutagenic primers by the AgeI site (5’ AAAAAAAAAACCGGTTTCTGCAGATATCCATCACACTGGCGGCCG 3’ and 5’ CGGCCGCCAGTGTGATGGATATCTGCAGAAACCGGTTTTTTTTTT 3’). The pcDNA3-FLUC plasmid containing the coding sequence of Firefly luciferase was a gift from Prof. Nahum Sonenberg^49^. The plasmid encoding yeast RPL41A was a gift from Prof. Jon Lorsch^50^. All plasmids were propagated in *E. coli* strain TOP10F’ competent cells.

#### Construction of GAPDH variants

To obtain the construct containing 53 CA repeats in the 5’ UTR of the GAPDH mRNA [abbreviated: GAPDH_5’ UTR (CA)_53_], the T7 promoter and 106 bp in the 5’ end of the 5’ UTR were excised from the GAPDH-encoding plasmid via digestion with BamHI and Bst36I and replaced with a fragment listed in Table 2 (GAPDH_5’ UTR (CA)_53_), which contained the T7 promoter sequence and 53 CA repeats. The fragment was PCR amplified using forward (5’ GTCTGTGGATCCTTCTAATACG 3’) and reverse (5’ACAGACCTGAGGTGTGTG 3’) primers and digested using BamHI and Bst36I. The construct containing 53 CA repeats in the GAPDH 3’ UTR [GAPDH_3’ UTR (CA)_53_] was generated by replacing the 152 bp KpnI-SacI sequence in the 3’ end of the 3’ UTR with a fragment containing the original 46 bp sequence downstream of the KpnI site and 53 CA repeats (indicated in Table 2). The latter fragment was PCR amplified using forward (5’ AAGAGAGGTACCCTCACTGCT 3’) and reverse (5’GGAAACAGCTATGAGAGCTC 3’) primers and digested using KpnI and SacI. The GAPDH constructs containing randomized or genetic sequences [GAPDH_3’ UTR shuffle; GAPDH_3’ UTR Genetic] in the GAPDH 3’ UTR were generated as described above by replacing the 152 bp KpnI-SacI sequence at the 3’ UTR of GAPDH with the fragments indicated in Table 2.

#### mRNA preparation

To prepare mRNAs, we employed run-off *in vitro* transcription catalyzed by 6-His-tagged T7 RNA polymerase. With the exception of the pcDNA3-FLUC plasmid containing the T3 promoter for transcription, all mRNA encoding sequences were cloned downstream of a T7 promoter. To obtain the transcripts with or without a 30-nt poly(A) tail, plasmid DNAs were digested by EcoRI or SacI, respectively, and precipitated with ethanol. To obtain the rabbit β-globin transcript with or without a 30-nt poly(A) tail, plasmid DNAs were digested by AgeI or SacI, respectively. 10 µg of linearized plasmid DNA was added to a 1 mL transcription reaction mixture containing 80 mM HEPES-KOH pH 7.5, 2 mM spermidine, 30 mM dithiothreitol, 25 mM NaCl, 8 mM MgCl_2_, and 0.8 mM each of ATP, UTP, GTP and CTP. The reaction was initiated by adding homemade T7 RNA polymerase to a final concentration of 2 µM and then incubated at 37°C for 4-6 hours. After the transcription reaction, the synthesized RNA was precipitated with 0.3 M sodium acetate pH 5.3 and 2.5 volumes of ethanol. The precipitated RNA was purified from a denaturing RNA gel (20 cm × 16 cm × 1.5 mm) in 7 M Urea, 1× TBE, and 5% acrylamide. The gel was pre-run at 20 mA for 30 min. An equal volume of formamide was added to the RNA before electrophoresis to aid denaturation of the RNA sample. RNA was run on the gel at 20 mA for at least 2 hours until the tracking dye, Bromophenol blue, migrated three-fourths of the way through the gel. RNA bands were visualized on a TLC plate by briefly exposing the gel to short wave (254 nm) UV light. The RNA was eluted/recovered from the gel slab in gel extraction buffer (0.3 M sodium acetate pH 5.3, 0.5% SDS, and 5 mM EDTA) followed by phenol-chloroform extraction and ethanol precipitation.

#### mRNA labeling with Cy3 and Cy5 fluorophores

The 5’ phosphate of the RNA was labeled using cystamine and a maleimide derivative of Cy5 (or Cy3, Click Chemistry) in the following steps as previously described^51^: (i) the 5’ γ phosphate of RNA was reacted with 1-Ethyl-3-[3-dimethylaminopropyl]carbodiimide hydrochloride (EDC, Thermo Scientific) and imidazole; (ii) the phosphorimidazolide derivative of RNA was reacted with cystamine; (iii) the product was reduced with TCEP to release the thiophosphate group, which was subsequently modified by Cy5 (or Cy3) maleimide for 2 hours at room temperature (RT). Labeled RNA molecules were purified using a 1 mL G-25 spin column equilibrated with ddH_2_O. We can reproducibly achieve 100% yield of 5’ phosphate labeling. The 5’-labeled RNAs were then subjected to 3’ end labeling. The RNA 3’ OH group was conjugated to pCp-Cy3 (or Cy5, Jena Bioscience) by T4 RNA ligase I in 20 mM MgCl_2_, 3.3 mM DTT, and 50 mM Tris-HCl pH 8.5 in the presence of 5 mM ATP and 10% DMSO. The efficiency of Cy3/5-pCp labeling of transcripts typically does not exceed 15-30%. To perform ensemble FRET experiments, the 5’ end of mRNAs was labeled with Cy3 to ensure that all acceptor (Cy5)-labeled RNAs were also labeled with a donor fluorophore. To perform single-molecule FRET experiments, the 5’ end of mRNAs was labeled with Cy5 to ensure that all donor (Cy3)-labeled RNAs, imaged using the green (532 nm) laser, were also labeled with an acceptor (Cy5) fluorophore. All labeled RNA molecules were gel purified by denaturing PAGE as described above and stored in ddH_2_O after desalting via a 1 mL G-25 spin column.

#### RNA folding

To measure the end-to-end distances of mRNAs by ensemble FRET, 300 nM doubly-labeled (5’-Cy3, 3’-Cy5) RNA samples were refolded in 30 μl of folding buffer (50 mM HEPES-KOH pH 7.5 and 100 mM KCl) by first heating RNA at 90 °C for 2 min and slowly cooling to 37 °C. At 37 °C, MgCl_2_ was added to a final concentration of 0.5 to 8 mM depending on the experiment and cooled to RT for 5 min. In most experiments, folding buffer contained 1 mM MgCl_2_. To disrupt the basepairing interactions between the 5’ and 3’ UTR in GAPDH or β-globin mRNAs (shown in Fig. 2), a 50-nucleotide long DNA oligomer complementary to the 3’ end of the 3’ UTR of mRNA was added during mRNA refolding (GAPDH: 5’ CCTGGTTGAGCACAGGGTACTTTATTGATGGTACATGACAAGGTGCGGCT 3’ and β-globin: 5’ CGCAATGAAAATAAATTTCCTTTATTAGCCAGAAGTCAGATGCTGAAGGG 3’).

For single-molecule experiments, 100 nM doubly-labeled RNA samples (5’-Cy5, 3’-Cy3) and 150 nM biotinylated anchor DNA oligomers in 5 μl folding buffer were annealed as described above. Anchor DNA oligomers were designed to have a minimal effect on the overall secondary structure using *OligoWalk* software^19^: GAPDH: 5’-biotin/GATGATCTTGAGGCTGTTG 3’; β-globin: 5’-biotin/TAGGATTGTTCATAACAGCA 3’; and MIF: 5’-biotin/CATGTCGTAATAGTTGATGT 3’.

#### Ensemble FRET measurements

The fluorescence emission of Cy3 and Cy5 were measured using a FluoroMax-4 (Horiba) spectrofluorometer at RT. A 12.5 mm × 45 mm quartz cuvette with a 10 mm path length (Starna Cells) was used for a sample volume of 30 µl. All measurements were performed in folding buffer (50 mM HEPES-KOH pH 7.5 and 100 mM KCl) in the presence of 0.5 to 8 mM MgCl_2_ depending on the experiment. Cy3 emission spectra (555-800 nm) were taken by exciting fluorescence at 540 nm. Cy5 emission spectra (645-800 nm) were taken by exciting fluorescence at 635 nm. The slit-widths for excitation and emission were set to 5 nm of spectral bandwidth. FRET efficiencies (*E*) between the 5’ and 3’ end-labeled mRNAs were determined from Cy3 and Cy5 emission spectra using the ratioA method^52^. RatioA for each experiment was calculated from the ratio of the extracted integrated intensity of the acceptor (Cy5) fluorescence, which is excited both directly by 540 nm light and by energy transfer, divided by the integrated intensity of the acceptor excited directly by 635 nm light. FRET efficiency (*E*) was determined according to the following equation:

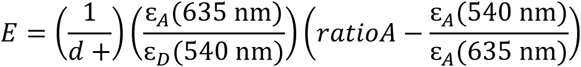

where d^+^ is the fraction of molecules labeled with donor. The donor labeling efficiency was determined from absorbance spectra. 1/d^+^ indicates the fraction of acceptor-labeled RNA that is also labeled with donor by assuming that the labeling of each fluorophore was random. ε is the extinction coefficient at the indicated wavelength, and the subscripts D and A denote donor or acceptor. The distance between donor and acceptor (R) was calculated from experimentally determined *E* values using the equation below:

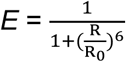

where R is the inter-dye distance and R_0_ is the Förster radius at which *E* = 0.5. Calculations of R were performed assuming R_0_= 56 Å^53^.

#### Single-molecule FRET measurements

100 nM biotin-conjugated Cy3/Cy5 doubly-labeled RNA samples were diluted in imaging buffer (50 mM HEPES pH 7.5, 100 mM KCl, 1mM MgCl_2_, 0.625% glucose, and 1.5 mM Trolox) to a final concentration of 50 pM and immobilized on quartz slides coated with biotinylated BSA (0.2 mg/mL, Sigma) and pre-treated with NeutrAvidin (0.2 mg/mL, Thermo Scientific). Imaging buffer with an oxygen-scavenging system (0.8 mg/mL glucose oxidase and 0.02 mg/mL catalase) was injected into the slide chambers before imaging to prevent photo-bleaching.

smFRET traces were recorded using a prism-based total internal reflection fluorescence (TIRF) microscope as previously described^54,55^. The flow chamber was imaged using an Olympus IX71 inverted microscope equipped with a 532 nm laser (Spectra-Physics) and 642 nm laser (Spectra-Physics) for Cy3 and Cy5 excitation, respectively. Fluorescence emission was collected by a water immersion objective (60×/1.20 w, Olympus). Fluorescence signals were split into Cy3 and Cy5 channels by a 630 nm dichroic beam splitter and recorded by EMCCD camera (iXon^+^, Andor Technology). Movies were recorded using Single software (downloaded from Prof. Taekjip Ha’s laboratory website at the Center for the Physics of Living Cells, University of Illinois at Urbana-Champaign (https://cplc.illinois.edu/software/)^12^ with the time resolution of 0.1 s for 10 min. All experiments were performed at RT.

#### Single-molecule data analysis

Collected datasets were processed and trajectories for individual molecules were extracted with IDL, using scripts downloaded from https://cplc.illinois.edu/software/. Apparent FRET efficiencies (*E*_app_) were calculated from the emission intensities of donor (*I*_Cy3_) and acceptor (*I*_Cy5_) as follows: *E*_app_ = *I*_Cy5_/ [*I*_Cy5_ + *I*_Cy3_]. The FRET distribution histograms were built from more than 200 trajectories that showed single-step disappearance for both Cy3 and Cy5 fluorescence intensities using a Matlab script provided by Prof. Peter Cornish (University of Missouri, Columbia). Single-step photobleaching of the acceptor dye resulting in a reciprocal increase in donor fluorescence indicated the presence of an energy transfer before acceptor photobleaching. FRET histograms were fitted to Gaussians using Origin (OriginLab). Among all collected traces, 20% of β-globin and GAPDH individual smFRET traces showed spontaneous interconversions between multiple FRET states. The state-to-state transitions in each fluctuating trace in GAPDH mRNA (N= 266) were determined using hidden Markov modeling (HMM) via HaMMy software^20^. smFRET traces showing apparent fluctuation were fit using HMM to 2, 3, and 4-state models. 99% of smFRET traces were best fit by 2-state models. Idealized FRET traces obtained by HMM were examined using transition density plot (TDP) analysis. To obtain TDP, the range of FRET efficiencies from 0 to 1 was separated into 200 bins. The resulting TDP heat map was normalized to the most populated bin in the plot. The lower- and upper-bound thresholds were set to 5% and 100% of the most populated bin, respectively.

### Computational procedures

#### Estimating end-to-end distance

To estimate end-to-end distance distributions and mean end-to-end distances for each RNA sequence, we used a two-scale freely jointed chain approximation^18^ for each structure in a Boltzmann ensemble of structures. We generate 1000 structures using stochastic sampling^24^ (program stochastic) in RNAstructure^23^. In stochastic sampling, structures are selected at random with the probability equal to their Boltzmann probability. Because the sample is Boltzmann weighted, the mean of a quantity across the sample has the proper Boltzmann weighting. For each structure, we count the number of branches and unpaired nucleotides in the exterior loop, i.e. the loop that contains the 5’ and 3’ ends, and use:

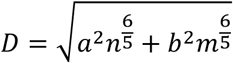

where *D* is the end-to-end distance, *n* is the number of unpaired nucleotides, *m* is the number of helical branches, *a* = 6.2 Å, and *b* = 15 Å, where a and b were from a previous parameterization^18^. The mean end to end distance is the arithmetic mean across structures in the sample.

#### Sequence Complexity

Sequence complexity is a measure of diversity for the nucleotide content of a sequence. In this work, we use *Linguistic complexity* as introduced by Trifonov^56^, and calculated using the algorithm from Gabrielian et al^26^. The complexity is the product of vocabulary size across k-mers:

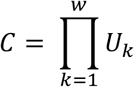

where the vocabulary size, *U*, is the fraction of possible sequences observed for that k-mer. The number of possible sequences for a k-mer is the minimum of 4^k^ or N-k+1, where N is the sequence length. For example, *k* = 3 has a possible sequence space of 4^3^ for sequences of 64 or more nucleotides, and *U*_3_ is the fraction of these 3-mer sequences observed across the sequence. The maximum k-mer size, *w*, is a function of length. Here we used *w* = 5 for the 106 nucleotide region of GAPDH mRNA and *w* = 7 for the full length GAPDH mRNA, following Gabrielian et al^26^.

#### Genetic Algorithm

We developed a genetic algorithm program to optimize features in a given RNA sequence. In this work, our goal was to evolve sequences to increase the end-to-end distance of the input sequences.

The genetic algorithm is an iterative process inspired by evolution in which an initial population is evolved to optimize features represented in the objective function^57^. A population of 10 sequences was used in this work, and these ten sequences were initialized uniformly as the starting sequence. In each iteration, sequences in the population are either mutated or new sequences are generated by recombining two sequences (called crossover) to generate 10 new sequences. The optimal 10 sequences (from the set of 10 at the start of the iteration and the 10 new sequences) are kept for subsequent iterations, where optimality is defined as maximizing the value of the objective function. In the mutation steps, each of the ten sequences was mutated independently. Sweeping along the portion of the sequence that is being evolved, there is a probability of 0.03 that a nucleotide will mutate to equal probability of A, C, G, or U. In our algorithm, crossover occurred every 6 steps. For crossover, 5 pairs of sequences are selected at random without replacement from the population of 10 sequences. For each sequence pair, the algorithm scans through the portion of the sequence that is being evolved and each nucleotide position has a probability of 0.03 to be selected as a recombination marker; therefore, on average, the number of recombination markers is 0.03×N. Then, the pair of sequences is recombined by the exchange of homologous segments to make two new sequences. The generation of the two sequences by crossover from the sequence pair is illustrated by the schematics shown below.

We used two objective functions in this work. In calculations shown in **Fig. 4**, the objective function was the mean probability of each nucleotide being unpaired as determined with a partition function calculation^58^. The mean is taken across only nucleotides that are in the region of the sequence being evolved. In calculations shown in **Fig. 5**, we summed the mean probability of each nucleotide being unpaired and the sequence complexity.

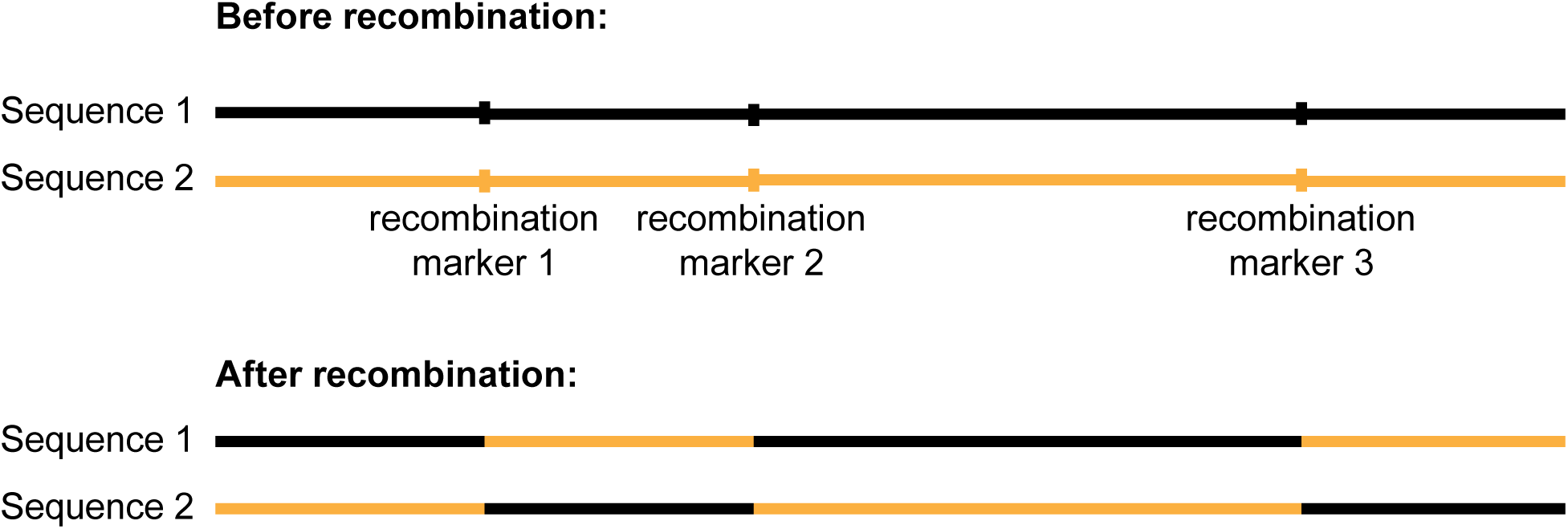

#### Genetic Algorithm Crossover

In crossover, two new sequences are generated from two sequences in the population. The black and orange colors indicate the sequence origin and final sequences. Recombination markers are chosen at random with frequency of 0.03 per nucleotide.

## ACKNOWLEDGEMENTS

This study was supported by grant from the US National Institute of Health GM-099719 (to D.N.E.) and R01GM076485 (to D.H.M). We thank Gloria Culver, Enea Salsi and Michael Sloma for their early contributions to this project.

## AUTHOR CONTRIBUTIONS

D.N.E. conceived the project. W-J.C.L., M.K., D.H.M. and D.N.E. designed research. W-J.C.L. performed experiments with contributions from E.V.C., E.F. and R.R. M.K. performed computational studies with contributions from S.B. W-J.C.L., M.K., E.V.C., D.H.M. and D.N.E. wrote the paper.

## REFERENCES

1. Hinnebusch, A.G., Ivanov, I.P. & Sonenberg, N. Translational control by 5’-untranslated regions of eukaryotic mRNAs. Science 352, 1413–6 (2016).

2. Gallie, D.R. The cap and poly(A) tail function synergistically to regulate mRNA translational efficiency. Genes Dev 5, 2108–16 (1991).

3. Tarun, S.Z., Jr. & Sachs, A.B. Association of the yeast poly(A) tail binding protein with translation initiation factor eIF-4G. EMBO J 15, 7168–77 (1996).

4. Tarun, S.Z., Jr., Wells, S.E., Deardorff, J.A. & Sachs, A.B. Translation initiation factor eIF4G mediates in vitro poly(A) tail-dependent translation. Proc Natl Acad Sci U S A 94, 9046–51 (1997).

5. Wells, S.E., Hillner, P.E., Vale, R.D. & Sachs, A.B. Circularization of mRNA by eukaryotic translation initiation factors. Mol Cell 2, 135–40 (1998).

6. Amrani, N., Ghosh, S., Mangus, D.A. & Jacobson, A. Translation factors promote the formation of two states of the closed-loop mRNP. Nature 453, 1276–80 (2008).

7. Sonenberg, N. & Hinnebusch, A.G. Regulation of translation initiation in eukaryotes: mechanisms and biological targets. Cell 136, 731–45 (2009).

8. Yoffe, A.M., Prinsen, P., Gelbart, W.M. & Ben-Shaul, A. The ends of a large RNA molecule are necessarily close. Nucleic Acids Res 39, 292–9 (2011).

9. Fang, L.T. The end-to-end distance of RNA as a randomly self-paired polymer. J Theor Biol 280, 101–7 (2011).

10. Clote, P., Ponty, Y. & Steyaert, J.M. Expected distance between terminal nucleotides of RNA secondary structures. J Math Biol 65, 581–99 (2012).

11. Leija-Martinez, N., Casas-Flores, S., Cadena-Nava, R.D., Roca, J.A., Mendez-Cabanas, J.A., Gomez, E. & Ruiz-Garcia, J. The separation between the 5’-3’ ends in long RNA molecules is short and nearly constant. Nucleic Acids Res 42, 13963–8 (2014).

12. Roy, R., Hohng, S. & Ha, T. A practical guide to single-molecule FRET. Nat Methods 5, 507–16 (2008).

13. Acker, M.G., Kolitz, S.E., Mitchell, S.F., Nanda, J.S. & Lorsch, J.R. Reconstitution of yeast translation initiation. Methods Enzymol 430, 111–45 (2007).

14. Pisarev, A.V., Unbehaun, A., Hellen, C.U. & Pestova, T.V. Assembly and analysis of eukaryotic translation initiation complexes. Methods Enzymol 430, 147–77 (2007).

15. Romani, A.M. Cellular magnesium homeostasis. Arch Biochem Biophys 512, 1–23 (2011).

16. Grosberg, A.Y. & Khokhlov, A.R. Statistical Physics of Macromolecules 350 (1994).

17. Cantor, C.R. & Schimmel, P.R. Biophysical Chemistry: Part I: The Conformation of Biological Macromolecules, 365 (1980).

18. Aalberts, D.P. & Nandagopal, N. A two-length-scale polymer theory for RNA loop free energies and helix stacking. RNA 16, 1350–5 (2010).

19. Mathews, D.H., Burkard, M.E., Freier, S.M., Wyatt, J.R. & Turner, D.H. Predicting oligonucleotide affinity to nucleic acid targets. RNA 5, 1458–69 (1999).

20. McKinney, S.A., Joo, C. & Ha, T. Analysis of single-molecule FRET trajectories using hidden Markov modeling. Biophys J 91, 1941–51 (2006).

21. Kinz-Thompson, C.D., Bailey, N.A. & Gonzalez, R.L., Jr. Precisely and Accurately Inferring Single-Molecule Rate Constants. Methods Enzymol 581, 187–225 (2016).

22. Furtig, B., Wenter, P., Reymond, L., Richter, C., Pitsch, S. & Schwalbe, H. Conformational dynamics of bistable RNAs studied by time-resolved NMR spectroscopy. J Am Chem Soc 129, 16222–9 (2007).

23. Reuter, J.S. & Mathews, D.H. RNAstructure: software for RNA secondary structure prediction and analysis. BMC Bioinformatics 11, 129 (2010).

24. 81 Ding, Y. & Lawrence, C.E. A statistical sampling algorithm for RNA secondary structure prediction. Nucleic Acids Res 31, 7280–301 (2003).

25. Majumdar, Z.K., Hickerson, R., Noller, H.F. & Clegg, R.M. Measurements of internal distance changes of the 30S ribosome using FRET with multiple donor-acceptor pairs: quantitative spectroscopic methods. J Mol Biol 351, 1123–45 (2005).

26. Gabrielian, A. & Bolshoy, A. Sequence complexity and DNA curvature. Comput Chem 23, 263–74 (1999).

27. Troyanskaya, O.G., Arbell, O., Koren, Y., Landau, G.M. & Bolshoy, A. Sequence complexity profiles of prokaryotic genomic sequences: a fast algorithm for calculating linguistic complexity. Bioinformatics 18, 679–88 (2002).

28. Varani, G. & McClain, W.H. The G x U wobble base pair. A fundamental building block of RNA structure crucial to RNA function in diverse biological systems. EMBO Rep 1, 18–23 (2000).

29. Chen, J.L., Dishler, A.L., Kennedy, S.D., Yildirim, I., Liu, B., Turner, D.H. & Serra, M.J. Testing the nearest neighbor model for canonical RNA base pairs: revision of GU parameters. Biochemistry 51, 3508–22 (2012).

30. Bichara, M., Pinet, I., Schumacher, S. & Fuchs, R.P. Mechanisms of dinucleotide repeat instability in Escherichia coli. Genetics 154, 533–42 (2000).

31. Rouskin, S., Zubradt, M., Washietl, S., Kellis, M. & Weissman, J.S. Genome-wide probing of RNA structure reveals active unfolding of mRNA structures in vivo. Nature 505, 701–5 (2014).

32. Wu, X. & Bartel, D.P. Widespread Influence of 3’-End Structures on Mammalian mRNA Processing and Stability. Cell 169, 905–917 e11 (2017).

33. Takyar, S., Hickerson, R.P. & Noller, H.F. mRNA helicase activity of the ribosome. Cell 120, 49–58 (2005).

34. Wen, J.D., Lancaster, L., Hodges, C., Zeri, A.C., Yoshimura, S.H., Noller, H.F., Bustamante, C. & Tinoco, I. Following translation by single ribosomes one codon at a time. Nature 452, 598–603 (2008).

35. Ding, Y., Tang, Y., Kwok, C.K., Zhang, Y., Bevilacqua, P.C. & Assmann, S.M. In vivo genome-wide profiling of RNA secondary structure reveals novel regulatory features. Nature 505, 696–700 (2014).

36. Aw, J.G. et al. In Vivo Mapping of Eukaryotic RNA Interactomes Reveals Principles of Higher-Order Organization and Regulation. Mol Cell 62, 603–17 (2016).

37. Lu, Z. et al. RNA Duplex Map in Living Cells Reveals Higher-Order Transcriptome Structure. Cell 165, 1267–1279 (2016).

38. Sharma, E., Sterne-Weiler, T., O’Hanlon, D. & Blencowe, B.J. Global Mapping of Human RNA-RNA Interactions. Mol Cell 62, 618–26 (2016).

39. Roth, A. & Breaker, R.R. The structural and functional diversity of metabolite-binding riboswitches. Annu Rev Biochem 78, 305–34 (2009).

40. Giedroc, D.P. & Cornish, P.V. Frameshifting RNA pseudoknots: structure and mechanism. Virus Res 139, 193–208 (2009).

41. Mauger, D.M., Siegfried, N.A. & Weeks, K.M. The genetic code as expressed through relationships between mRNA structure and protein function. FEBS Lett 587, 1180–8 (2013).

42. Leipuviene, R. & Theil, E.C. The family of iron responsive RNA structures regulated by changes in cellular iron and oxygen. Cell Mol Life Sci 64, 2945–55 (2007).

43. Simon, A.E. & Miller, W.A. 3’ cap-independent translation enhancers of plant viruses. Annu Rev Microbiol 67, 21–42 (2013).

44. Li, X., Quon, G., Lipshitz, H.D. & Morris, Q. Predicting in vivo binding sites of RNA-binding proteins using mRNA secondary structure. RNA 16, 1096–107 (2010).

45. 102 Li, X., Kazan, H., Lipshitz, H.D. & Morris, Q.D. Finding the target sites of RNA-sbinding proteins. Wiley Interdiscip Rev RNA 5, 111–30 (2014).

46. Archer, S.K., Shirokikh, N.E., Hallwirth, C.V., Beilharz, T.H. & Preiss, T. Probing the closed-loop model of mRNA translation in living cells. RNA Biol 12, 248–54 (2015).

47. Babendure, J.R., Babendure, J.L., Ding, J.H. & Tsien, R.Y. Control of mammalian translation by mRNA structure near caps. RNA 12, 851–61 (2006).

48. Kozak, M. Influences of mRNA secondary structure on initiation by eukaryotic ribosomes. Proc Natl Acad Sci U S A 83, 2850–4 (1986).

49. Poulin, F., Gingras, A.C., Olsen, H., Chevalier, S. & Sonenberg, N. 4E-BP3, a new member of the eukaryotic initiation factor 4E-binding protein family. J Biol Chem 273, 14002–7 (1998).

50. Mitchell, S.F., Walker, S.E., Algire, M.A., Park, E.H., Hinnebusch, A.G. & Lorsch, J.R. The 5’-7-methylguanosine cap on eukaryotic mRNAs serves both to stimulate canonical translation initiation and to block an alternative pathway. Mol Cell 39, 950–62 (2010).

51. Lai, W.C. & Ermolenko, D.N. Ensemble and single-molecule FRET studies of protein synthesis. Methods (2017).

52. Clegg, R.M. Fluorescence resonance energy transfer and nucleic acids. Methods Enzymol 211, 353–88 (1992).

53. Dietrich, A., Buschmann, V., Muller, C. & Sauer, M. Fluorescence resonance energy transfer (FRET) and competing processes in donor-acceptor substituted DNA strands: a comparative study of ensemble and single-molecule data. J Biotechnol 82, 211–31 (2002).

54. Cornish, P.V., Ermolenko, D.N., Noller, H.F. & Ha, T. Spontaneous intersubunit rotation in single ribosomes. Mol Cell 30, 578–88 (2008).

55. Salsi, E., Farah, E., Dann, J. & Ermolenko, D.N. Following movement of domain IV of elongation factor G during ribosomal translocation. Proc Natl Acad Sci U S A 111, 15060–5 (2014).

56. Trifonov, E.N. Making sense of the human genome. in Structure and methods, Vol. 1 (eds. Sarma, R.H. & Sarma, M.H.) 69-77 (Adenine Press, 1990).

57. Forrest, S. Genetic algorithms: principles of natural selection applied to computation. Science 261, 872–8 (1993).

58. Mathews, D.H. Using an RNA secondary structure partition function to determine confidence in base pairs predicted by free energy minimization. RNA 10, 1178–90 (2004).

