## Supplementary Materials for "The formation of intramolecular secondary structure brings mRNA ends in close proximity"

### Supplementary results:

**Supplementary Table 1. FRET-derived and computationally predicted end-to-end distances in various natural mRNAs and GAPDH mRNA variants.**

| mRNA | Species | Length (nt) | FRET derived end-to-end distance (nm) | Predicted end-to-end distance (nm) |
|---|---|---|---|---|
| F-Luciferase (ORF) | Firefly | 1689 | 4.9±0.1 | 3.5 |
| β-globin | Rabbit | 597 | 4.7±0.1 | 4.6 |
| Ribosomal protein L41A (RPL41A) | Yeast | 321 | 6.7±0.2 | 3.5 |
| Heat shock binding protein 1 (HSBP1) | Human | 533 | 6.6±0.2 | 2.7 |
| ATP synthase subunit F (ATP5J2) | Human | 443 | 6.7±0.5 | 2.7 |
| Macrophage migration inhibitory factor (MIF) | Human | 561 | 4.8±0.3 | 3.1 |
| Mitochondrial ribosomal protein L51 (MRPL51) | Human | 690 | 4.3±0.4 | 2.6 |
| Glyceraldehyde-3-phosphate dehydrogenase (GAPDH) | Human | 1327 | 6.0±0.1 | 2.6 |
| GAPDH 3' UTR shuffle | Human | 1327 | 6.1±0.2 | 2.5 |
| β-globin_polyA$_{30}$ | Rabbit | 627 | 9.9±0.4 | 6.5 |
| GAPDH_polyA$_{30}$ | Human | 1357 | 11.2±0.5 | 6.2 |
| GAPDH 5' UTR(CA)$_{53}$ | Human | 1327 | n.d. | 11 |
| GAPDH 3' UTR(CA)$_{53}$ | Human | 1327 | n.d. | 11 |
| GAPDH 3’ UTR genetic | Human | 1327 | 9.4±0.3 | 10.4 |

Average FRET-derived end-to-end distances and respective SD values were determined from three to five independent mRNA refolding experiments performed in the presence of 1 mM $MgCl_2$ and 100 mM KCl. The computationally predicted end-to-end distances represent the mean distance in the ensemble of 1,000 structures generated by stochastic sampling with the *RNAstructure* softare package for each mRNA.

**Supplementary Table 2. Rates of transitions between different FRET states in GAPDH mRNA.**

| Transition | Number of transitions | Rate, $s^{-1}$ |
|---|---|---|
| 0.4 → 0.6 | 2112 | 0.13 ± 0.02 |
| 0.6 → 0.4 | 2050 | 0.11 ± 0.02 |
| 0.6 → 0.8 | 492 | 0.14 ± 0.03 |
| 0.8 → 0.6 | 460 | 0.03 ± 0.01 |

Rates were determined from 5,114 fluctuations between different FRET states in 266 HMM-idealized FRET traces obtained for GAPDH mRNA folded in presence of 1 mM $MgCl_2$ and 100 mM KCl.

**Supplementary Figure 1:**

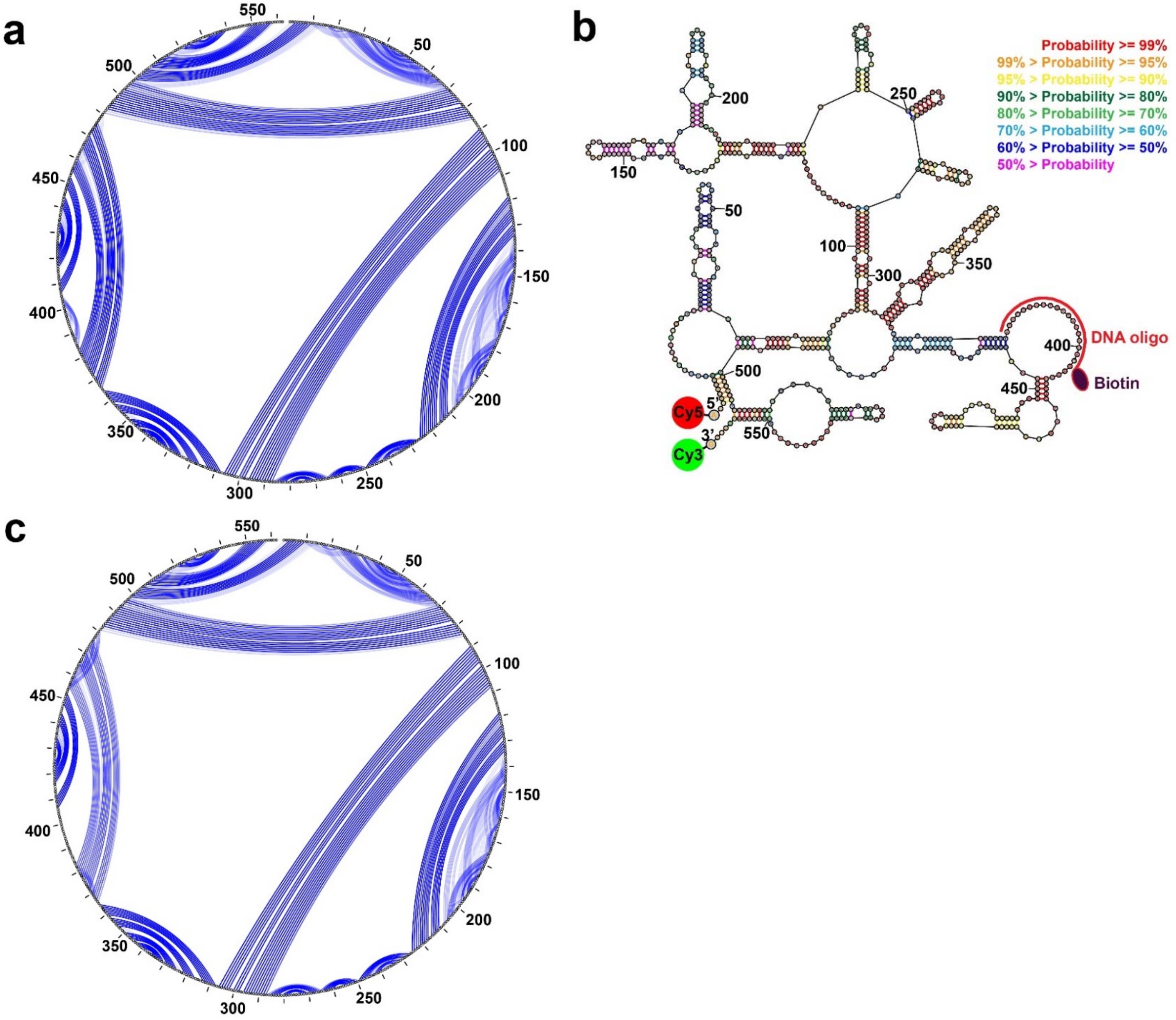


**Supplementary Figure 1. Secondary structures of human *MIF* mRNA lacking poly(A) tail predicted by free energy minimization. (a)** The circle diagram depicts base pairings, which are represented by the arcs, in the secondary structure of human *MIF* mRNA lacking poly(A) tail. This image is produced with the StructureEditor program from the RNAstructure software package (https://rna.urmc.rochester.edu/RNAstructure.html). **(b)** Secondary structure of human *MIF* mRNA lacking poly(A) tail folded in the presence of a 20-nucleotide biotinylated DNA oligomer. The biotinylated DNA oligomer, which was designed by *OligoWalk* to have a minimal impact on the mRNA structure, was used for the immobilization of fluorescently-labeled mRNA in smFRET experiments. The predicted lowest free energy structure was drawn with the draw program in *RNAstructure (https://rna.urmc.rochester.edu/RNAstructure.html)*. Base pair probabilities, predicted with a partition function, are indicated by the color key. In order to measure the end-to-end distance by FRET, the 5' and 3' ends of mRNA were conjugated with acceptor (red) and donor (green) fluorophores, respectively, as indicated. **(c)** The circle diagram depicts base pairings, which are represented by the arcs, in the structure shown in panel **b**.

**Supplementary Figure 2:**

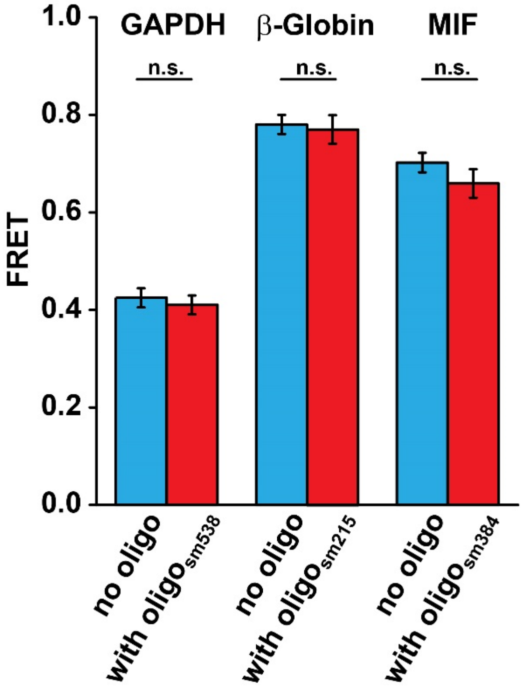


**Supplementary Figure 2. Annealing of biotin-labeled DNA oligomers does not affect end-to-end distance in GAPDH, β-globin and MIF mRNA.** Biotinylated DNA oligonucleotides (sm538 for GAPDH mRNA, sm215 for β-globin mRNA, and sm384 for MIF mRNA), which were predicted to have a minimal effect on the overall secondary structure and end-to-end distance, were used to tether the mRNAs to microscope slide in smFRET experiments shown in **Fig. 2**. FRET values measured in mRNAs folded in the presence (red) or in the absence (blue) of biotinylated DNA oligomers were not statistically significant (n.s.), as determined by the Student t-test (with α of 0.05). Each FRET value represents the mean ± SD of three independent experiments.

**Supplementary Figure 3:**

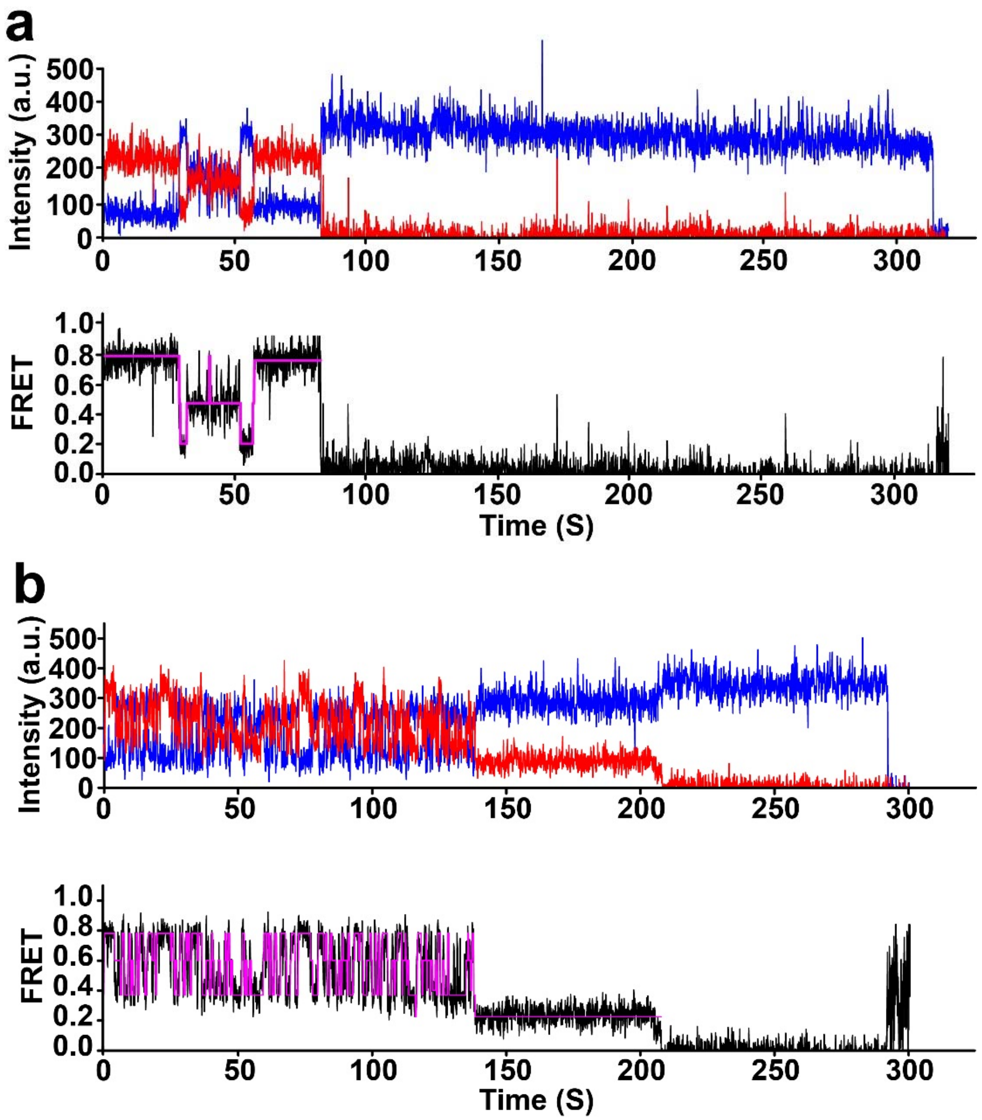


**Supplementary Figure 3. smFRET traces for GAPDH mRNA showing fluctuations between 0.2, 0.4 and 0.8 (panel a) or 0.2, 0.4, 0.6 and 0.8 FRET states (panel b).** Observed intensities of donor and acceptor fluorescence and the calculated apparent FRET efficiency are shown in blue, red and black, respectively. The Hidden Markov Model fit is shown in magenta.

**Supplementary Figure 4:**

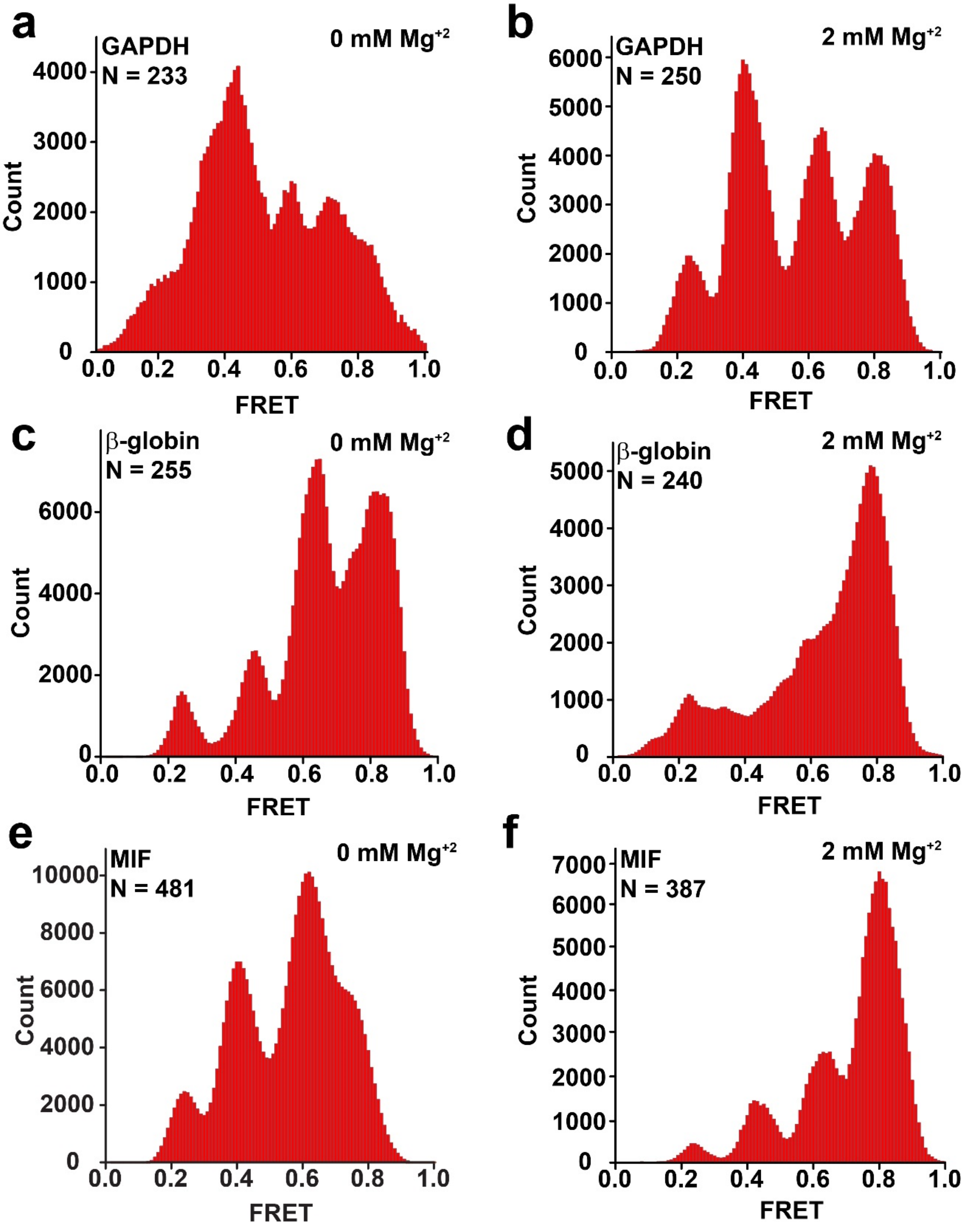


**Supplementary Figure 4. FRET distribution histograms for GAPDH (a-b), β-globin (c-d) and MIF (e-f) mRNAs folded and imaged in the absence of $MgCl_2$ (a, c, e) or in the presence of 2 mM $MgCl_2$ (b, d, f).** N is the number of traces used to assemble each histogram.

**Supplementary Figure 5:**

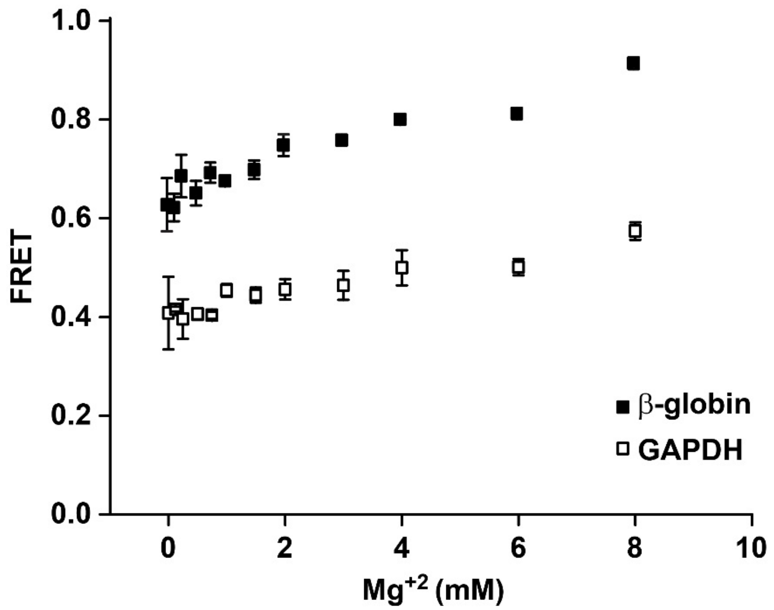


**Supplementary Figure 5. Efficiency of energy transfer between fluorophores attached to mRNA ends as a function of $MgCl_2$ concentration.** GAPDH (open squares) and β-globin (filled squares) mRNAs were folded in the presence of different $MgCl_2$ concentrations ranging from 0 to 8 mM. Error bars show standard deviations calculated from three to five independent experiments.

**Supplementary Figure 6:**

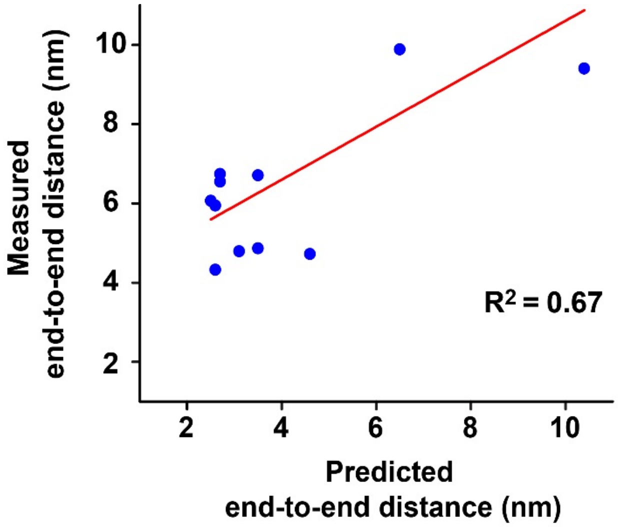


**Supplementary Figure 6. Correlation between computationally predicted (x-axis) and experimentally determined (y-axis) end-to-end distances.** End-to-end distance was predicted and measured by FRET in yeast RPL41A, firefly luciferase, rabbit β-globin, human ATP5J2, HSBP1, MIF, MRPL51 and GAPDH mRNAs, all of which lacked poly(A) tail. In addition, end-to-end distance was predicted and measured by FRET in rabbit β-globin and human GAPDH mRNAs containing a 30 nt-long poly(A) tail.
